# Detection of Domestication Signals through the Analysis of the Full Distribution of Fitness Effects

**DOI:** 10.1101/2022.08.24.505198

**Authors:** David Castellano, Ioanna-Theoni Vourlaki, Ryan N. Gutenkunst, Sebastian E. Ramos-Onsins

## Abstract

Domestication is a process marked by complex interactions between demographic changes and selective pressures, which together shape genetic diversity. While the phenotypic outcomes of domestication are well documented, its genetic basis—particularly the dynamics of selection— remain less well understood. To investigate these dynamics, we performed simulations designed to approximate the demographic history of large domestic mammals. These simulations used selection coefficients as a modeling tool to represent changes in selection pressures, recognizing that such coefficients are abstractions rather than direct representations of biological reality. Specifically, we analyzed site frequency spectra (SFS) under varying distributions of fitness effects (DFE) and proportions of mutations with divergent selective pressures. Our results show that the discretized deleterious DFE can be reliably inferred from the SFS of a single population, but reconstructing the beneficial DFE and demographic history remains challenging, even when using the joint SFS of both populations. We further developed a novel joint DFE inference model to estimate the proportion of mutations with divergent selection coefficients (*p*_c_), although we found that signals of classic hard sweeps can mimic increases in *p*_c_, complicating interpretation. These findings underscore both the utility and limitations of DFE inference and highlight the need for caution when interpreting demographic histories in domesticated populations based on such modeling assumptions.

## INTRODUCTION

The increase in human population and the emergence of modern society are closely linked to the domestication of plants and animals (Purugganan and Fuller 2009; Driscoll *et al*. 2009; Larson and Burger 2013; Amills *et al*. 2017). Human civilization was made possible through the domestication of surrounding life forms, where plants and animals such as wheat, dogs, pigs, or chickens were among the first to be domesticated (Dayan 1994; Zeder *et al*. 2006; Zeder 2012, 2015; Redding 2015; Avni *et al*. 2017). Domestication is a process that fosters long-term mutualistic relationship, providing benefits to both humans and domesticated species (Zeder *et al*. 2006). This process began approximately 10-15 thousand years ago and continues to this day (Larson and Burger 2013; Zeder 2015).

Despite its foundational role in human civilization, our genomic and evolutionary understanding of domestication remains incomplete. Domestication occurs rapidly on evolutionary time scale, but it is not a discrete event; rather, it involves the gradual refinement of domesticated traits. Artificial selection during domestication is often assumed to be relatively stronger and faster than natural selection. However, evidence from plants suggests that the evolutionary rate of domesticated varieties can be similar to that of wild plants, indicating a process more akin natural selection (Purugannan and Fuller 2010).

Domestication is also commonly associated with population bottlenecks; where only a small subset of individuals from a wild population are domesticated, potentially reducing the efficiency of natural selection (Wright *et al*. 2005). An additional distinction between natural and artificial selection is the use of truncation selection by modern breeders -a method that selects the top percentage of individuals for desired traits (Granleese *et al*. 2019). The prevalence of truncation selection in natural populations or prior to industrialization remains unknown. Truncation selection is an efficient form of directional selection (Crow and Kimura 1979), and significant genetic load accumulation is unlikely in outcrossing species (Kondrashov 1988; Ohta 1989) if the population sizes remain sufficiently large (Marsden *et al*. 2016).

A recent comprehensive meta-analysis of the genetic costs of domestication (Moyers *et al*. 2018) revealed that domesticated populations generally carry more deleterious variants, or segregate at higher frequencies, compared to their wild counterparts. However, this pattern is not universal, as evidenced by studies in sorghum (Lozano et al. 2021). Such patterns are likely driven by multiple processes that reduce the effectiveness of selection in domesticated populations, a concept first observed in rice genomes (Lu *et al*. 2006).

Selection, both natural and artificial, can act through a few loci with strong effects or many loci with small effects, depending on the genetic architecture of the trait and the strength of selection (Jain and Stephan 2017a; b). These two selection models are expected to produce distinct patterns of genetic diversity around selected loci (Stephan and John 2020). Classic hard selective sweeps have been reported at a few candidate loci for key domesticated traits (reviewed by Andersson 2012), such as the IGF2 gene region associated with lean pigs (Van Laere *et al*. 2003), the thyroid-stimulating hormone receptor (TSHR) in chickens (Rubin *et al*. 2010), and the sh4 and qSW5 loci related to seed shattering and grain width in rice (Shomura *et al*. 2008; Li *et al*. 2018; Huang *et al*. 2012). These cases reflect a Mendelian genetic architecture, where a small number of loci explain most of the phenotypic variance (see Courtier-Orgogozo and Martin 2020 for a comprehensive list of genes related to domestication). In short, genomic analyses of domestication have traditionally focused on identifying strong selection footprints, often driven by loci with large effects responsible for phenotypic differences (e.g. Groenen et al 2012; Carneiro et al 2019; Qanbari et al 2019; Li et al 2020). However, Leno-Colorado et al. (2022) found that domesticated and wild pig populations did not differ in the number and type of non-synonymous fixed mutations, contradicting the idea that most domesticated traits follow a Mendelian genetic architecture. Thus, the hard selective sweep model may be the exception rather than the rule in pigs domestication.

In this study, we investigate the genomic consequences of domestication by modeling and comparing the full distribution of fitness effects (DFE) for new and standing genetic variation. A change in the selection regime can be modeled in different ways: as shifts in selection coefficients, as done here, or alternatively, as changes in the optimal value of a quantitative trait determined by a set of loci whose effects on fitness depend on their contribution to the trait and the genetic background they are in. In our approach, we infer the joint DFE for wild and domesticated populations using selection coefficients as abstractions to approximate the effects of selection. This allows us to quantify the proportion of shared genetic variants (modeled as) having diverging selection pressures, providing insights into how selective regimes may differ between these populations. We recognize that alternative frameworks, such as quantitative trait models, may offer complementary perspectives on the genetic consequences of domestication.

Previous studies estimating the DFE have primarily relied on contrasting the site frequency spectrum (SFS) of synonymous and non-synonymous mutations within a single population (1D-SFS). These methods assume that beneficial mutations contribute to divergence but not to polymorphism due to their rapid fixation (Keightley and Eyre-Walker 2007; Boyko *et al*. 2009; Kim *et al*. 2017; Tataru *et al*. 2017; Barton and Zeng 2018). Tataru *et al*. (2017) developed polyDFE to infer the full DFE and the proportion of adaptive substitutions (α) using polymorphism data alone. They also proposed nested models to test whether the parameters of the DFEs are shared between populations. Castellano et al (2019) applied polyDFE to great apes and found that the shape parameter of the gamma deleterious DFE is likely conserved across these closely related species. However, populations that have diverged much more recently than great apes -such as domesticated and wild populations-tend to share a large number of genetic variants. To better leverage this shared variation, Huang et al. (2021) proposed using the SFS of both populations simultaneously (2D-SFS) to jointly estimate the deleterious DFE. Traditional 1D-SFS-based methods only provide access to the marginal DFE of the population without the need for shared variants, as they do not focus on the selection coefficients of individual mutations. In contrast, joint 2D inference, while more limited in applicability due to its reliance on substantial shared genetic variation, offers the advantage of quantifying the stability of the direction and intensity of natural selection on individual mutations.

Inferring the demographic history of domesticated populations is as important as inferring the change in the selection regime between domesticated and wild populations. Demographic processes associated with domestication have been studied across several species (Morell Miranda et al., 2023; Arnoux et al., 2020; Murray et al., 2010), with key events such as population splits between wild and domesticated groups, bottlenecks, and gene flow being inferred. These studies have also considered the influence of multiple selective sweeps (Caicedo et al., 2007). However, it has been demonstrated that ignoring background selection when analyzing demographic patterns can result in biased estimates (Torres et al., 2020; Comeron, 2017; Beissinger et al., 2016). We will revisit this issue and provide broader context in the results and discussion.

To gain insights into the inference of complex demographic histories and the DFE in the context of domestication, we employ forward-in-time simulations of an idealized domestication process. These simulations explore various demographic and selective scenarios, enabling us to evaluate the ability to detect differences in selective pressures between domesticated and wild populations. We simulate a range of genetic architectures and selective effects, including: (1) Models with a relatively small number of loci undergoing changes in their selective effects and (2) models where numerous loci exhibit divergent selective effects. We also play with the rate and mean effect of beneficial mutations to understand the role of selective sweeps. Importantly, we introduce a novel methodology based on Huang *et al*. (2021) that incorporates an additional parameter critical for distinguishing populations experiencing rapid selective change: the selective effects of a fraction of existing variants can change (e.g., from beneficial to deleterious, or vice versa) in domesticated populations. This method jointly infers the full DFE parameters for both wild and domesticated populations, including shifts in the selective effects of shared variants. Finally, we describe and discuss how linked selection and changes in the DFE impair our ability to accurately infer the true simulated demographic histories.

## MATERIALS AND METHODS

### Simulation of the Domestication Process

A simulation of an idealized domestication process is developed using the forward-in-time simulator SLiM4 (Haller and Messer 2023). The general model of the domestication process is developed in the SLiM script available in github (https://github.com/CastellanoED/domesticationDFE) and Zenodo (https://zenodo.org/records/14277802). Twenty-four different domestication scenarios are analyzed, and the parameters for each scenario are shown in Table 1. All options (flags) used to run the SLiM script are also available in github (https://github.com/CastellanoED/domesticationDFE/blob/main/slim_code_mod4_NEW.slim). We aim to model a general domestication process that resembles the genomic configuration, generation time, mutation and recombination landscape relevant to large domesticated mammals used as livestock. Note that we are not considering the recent processes of genetic improvement performed by commercial companies in the last decades. The constructed model assumes a genome containing a single chromosomal “chunk” or window, with 10,000 loci/exons of 120 base pairs in length, and each locus/exon with one-third (4-fold) neutral synonymous positions and two-thirds (0-fold) selected non-synonymous positions scattered along the locus.

**Table 1.** Variable demographic and selective parameters across scenarios.

| **Migration**<br>**(W->D)** | **Positive DFE** | **$p_c$** | **Scenario**<br>**ID** |
| --- | --- | --- | --- |
| 0 | $p_b = 0$ | 0 | 1 |
|  | Absent | 0.05 | 2 |
|  |  | 0.25 | 3 |
| | $p_b = 0.1$ & $S_b = 1$ | 0 | 4 |
|  | Pervasive and nearly neutral | 0.05 | 5 |
|  |  | 0.25 | 6 |
| | $p_b = 0.01$ & $S_b = 10$ | 0 | 7 |
|  | Common and weak | 0.05 | 8 |
|  |  | 0.25 | 9 |
| | $p_b = 0.001$ & $S_b = 100$ | 0 | 10 |
|  | Rare and strong | 0.05 | 11 |
|  |  | 0.25 | 12 |
| 0.01 | $p_b = 0$ | 0 | 13 |
|  | Absent | 0.05 | 14 |
|  |  | 0.25 | 15 |
| | $p_b = 0.1$ & $S_b = 1$ | 0 | 16 |
|  | Pervasive and nearly neutral | 0.05 | 17 |
|  |  | 0.25 | 18 |
| | $p_b = 0.01$ & $S_b = 10$ | 0 | 19 |
|  | Common and weak | 0.05 | 20 |
|  |  | 0.25 | 21 |
| | $p_b = 0.001$ & $S_b = 100$ | 0 | 22 |
|  | Rare and strong | 0.05 | 23 |
|  |  | 0.25 | 24 |

The twenty-four simulated combinations of parameters in this study. The first column refers to the DFE of new beneficial mutations, the second column represents the migration rate from the Wild to the Domesticated population and the third column shows the probabilities to have sites that change their selection coefficients in the Domesticated population ( $p_c$ ). Last column shows the ID we use to quickly label scenarios along the manuscript.

The simulation parameters for each scenario (Figure 1A) are as follows: the initial population at time 0 run for 10\**N*e generations to reach mutation-selection-drift equilibrium, then splits into the domesticated and wild populations. Hereafter we refer to the Wild and Domesticated populations. We aim to mimic a realistic but still general domestication process in large livestock mammals where ancestral Ne (N_a_) estimates are on average around 10,000 (Murray *et al*. 2010; Groenen *et al*. 2012; Larson *et al*. 2014; Frantz *et al*. 2015; Yang *et al*. 2016; Librado *et al*. 2021; Todd *et al*. 2022) and the domestication process, according to archeological records, started around 10,000 years ago (Ahmad *et al*. 2020). The average generation time in these large domesticated mammals is about 5 years per generation (Pacifici *et al*. 2013). Note that in this study we had to reduce the population size and related population parameters below from 10,000 diploid individuals to N_a_=5,000 diploid individuals for computational reasons.

**Figure 1:**
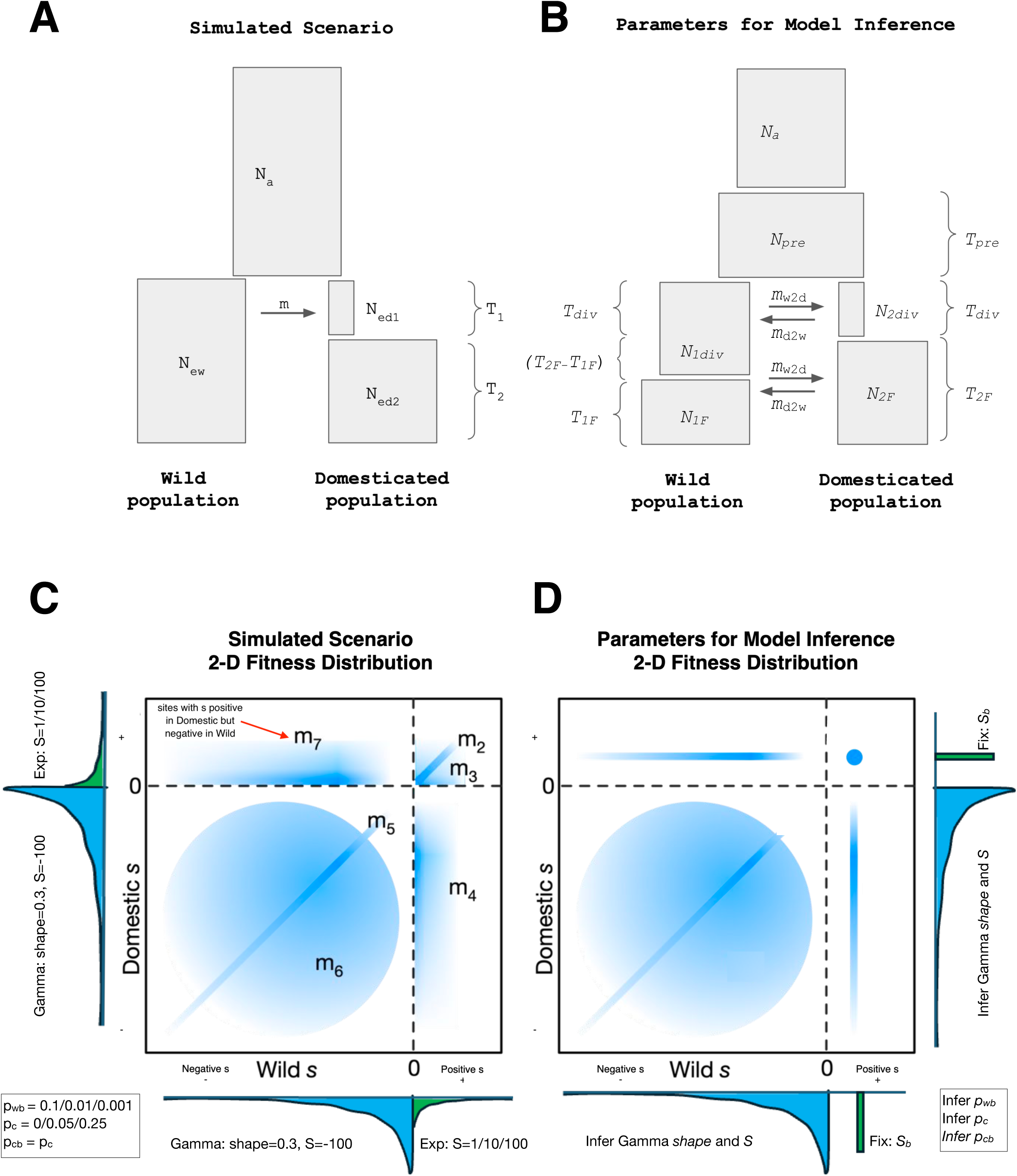
Joint demographic and DFE models simulated and fit. **A:** Illustration of the joint demographic model used in SLiM simulations. N_a_: Effective population size of the Ancestral population. N_ew_: Effective population size of the Wild population. N_e1d_: Effective population size of the Domesticated population during the bottleneck. N_e2d_: Effective population size of the Domesticated population after the bottleneck. T_1_: Number of generations in the bottleneck period. T_2_: Number of generations from the bottleneck to the present. m: Wild to Domesticated migration rate (migration occurs along T_1_). **B:** Illustration of a more general joint demographic model used in the dadi inferences. *N_a_*: Effective population size of the Ancestral population. *N_pre_*: Effective population size before the domestication split. *N_1div_*: Effective population size of the Wild population after the split. *N_1F_*: Effective population size of the Wild population at the end of the simulation. *N_2div_*: Effective population size of the Domesticated population after the split. *N_2F_*: Effective population size of the Domesticated population at the end of the simulation. T_pre_: Number of generations before the domestication split. *T_div_*: Number of generations after the bottleneck. *T_1F_*: Number of generations under *N_1F_*. *T_2F_*: Number of generations under *N_2F_*. Note that *T_1F_* and *T_2F_* are estimated independently and that *T_1F_* can be the same, longer or shorter than *T_2F_*. *m_d_*: Wild to Domesticated migration rate. *m_w_*: Domesticated to Wild migration rate. Both migration rates occur after the domestication split. **C:** Illustration of the joint DFE model used in the SLiM simulations, with mutation types illustrated. In the illustration, the shadow blue regions in the plot represent the possible different types of mutations considering the selection coefficient values in each of the two populations (from gamma and exponential distributions in wild and domestic and from the proportions of p_wb_, pc and p_cb_, see Table 1 and 2). For example, a point in the left-upper region of the illustration represents a mutation with positive s in the Domestic population but negative in Wild population (type m_7_). **D:** Illustration of the joint DFE model used in the dadi inferences and the inferred associated parameters, in which a fixed positive selection coefficient is assumed.

Genomic parameters: The mutation rate per site (μ) and generation is 2.5×10^−7^, and the population size (N_a_) is 5,000 diploid individuals, thus the expected θ under neutrality is 0.005. Each locus is separated from its neighbors by 3×10^−6^ recombination events per generation. The recombination rate per site and generation within the loci is fixed to a rate of 1.5×10^−7^ recombination events per site. Note that the higher recombination between loci aims to mimic their real genetic distance separation (assuming a functional site density of 5%) - this greatly speeds up the simulation as non-coding sites do not need to be simulated. In other words, we simulate 120 Kb of coding sequence in each run, which is equivalent to simulating a 2.4 Mb chromosome window with 5% coding sites. We perform 100 independent runs for each of the twenty-four scenarios.

Demographic parameters: The Domesticated populations of 5,000 diploid individuals suffer a bottleneck, reducing their population size temporarily to 200 diploid individuals, to recover again to 5,000 diploid individuals after the bottleneck. The bottleneck lasted 100 generations. The simulation finishes 900 generations after the bottleneck. In twelve of the twenty-four simulated scenarios we allow that a ratio of 0.01 of the Wild individuals migrate to the Domesticated population during the 100 generations of the bottleneck. Thus, during the bottleneck 25% of the domesticated population comes from the wild population every generation. In the other twelve combinations there is no exchange of individuals between the Wild and Domesticated populations. This demographic history is equivalent to a 1,000 years long bottleneck followed by a 9,000 years long recovery in an ancestral population with 10,000 diploid individuals and a generation time of 5 years.

DFE parameters: the selective effects produced by domestication are modeled by changing the fitness values of a proportion (calculated with a probability of change called p_c_) of the existing and new mutations in the domesticated population (at the time of the split) (Table 1). This probability of change can be 0 (our negative control), 0.05 or 0.25. Domesticated and Wild populations show different proportions of beneficial and deleterious new mutations depending on the scenario. SLiM defines ‘s*’* as the selective coefficient for the homozygote, while the inference algorithms used estimate the selective coefficient for heterozygote. Here we have assumed co-dominance and we have scaled the coefficient of selection to the ancestral population to have comparative values, that is, we multiply the N_a_ 4 times and divide the selective coefficient twice, 4N_a_s/2=2N_a_s). The negative effects in all scenarios and populations follow a gamma distribution with a shape value of 0.3 and a mean of S = −100 (S =2N_a_s in the heterozygote, in which each mutation is assumed co-dominant, N_a_ = 5,000 diploid individuals, considering the ancestral population size and s = −-0.01), which is in the range of values inferred using empirical data (Boyko et al. 2008, Galtier 2016), while variants with positive effects follow an exponential distribution. We investigate three combinations of parameters for the positive DFE plus one without positive selection: no positive selection (that is, p_b_ = 0), pervasive & nearly neutral (with mean selective effects S_b_ = 1 and probability of being beneficial p_b_ = 0.10), common & weak (S_b_ = 10 and p_b_ = 0.01) and rare & strong (S_b_ = 100 and p_b_ = 0.001) (Table 1).

### Types of Sites

The sites are initially divided into seven different types (named m_1_ to m7), being m_1_ neutral (synonymous) and m_2_ to m_7_ functional (non-synonymous) sites having a different selective effect when mutated (see Table 2 and Figure 1C). Mutations at m_5_, m_6_ and m_7_ sites generate deleterious variants in the Wild population, and mutations at m_2_, m_3_ and m_4_ sites generate beneficial mutations in the Wild population. The selection coefficient of mutations generated at m_2_ (beneficial) or at m_5_ (deleterious) sites are invariant for the Wild and Domesticated populations. However, the mutations at m_3_, m_4_, m_6_ and m_7_ sites will change their selective effect in the Domesticated populations relative to the Wild populations. That is, the new selective effect is drawn from the corresponding DFE section (positive or negative), independently of their value in the wild population. The selection coefficient of a given beneficial mutation at m_3_ sites will remain beneficial in the Domesticated population, but it will be different from the original beneficial effect at Wild. A mutation at m_4_ sites will change its selection coefficient from beneficial in the Wild to deleterious in the Domesticated population. Equivalently, the selection coefficient of a deleterious mutation at m_6_ sites will remain negative in the Domesticated population but it will be different from that found at Wild. A mutation at m_7_ sites will change its selection coefficient from deleterious in the Wild to beneficial in the Domesticated population (see probabilities included in Table 2). For each of the 24×100 simulation runs, we randomly pre-calculate independently the location of each site type (except for the permanent location of two non-synonymous sites followed by a synonymous site within codons) and their selective effect using an ad hoc R script (https://github.com/CastellanoED/domesticationDFE/blob/main/calculate_fitness_position_matrix.R). This hard-coding of selective effects on different sites allows us to gain insight into the relative importance of each mutation type for the domestication process.

**Table 2.** Types of sites in simulated scenarios.

| Site | Wild | Domesticated | Probability<sup>1</sup> |
| --- | --- | --- | --- |
| $m_1$ | Neutral | No change, remain Neutral | All synonymous |
| $m_2$ | Beneficial | No change, remain Beneficial | $p_{wb} \cdot (1 - p_c)$ |
| $m_3$ | Beneficial | Change to a different Beneficial Effect | $p_{wb} \cdot p_c \cdot p_{cb}$ |
| $m_4$ | Beneficial | Change to Deleterious | $p_{wb} \cdot p_c \cdot (1 - p_{cb})$ |
| $m_5$ | Deleterious | No change, remain Deleterious | $(1 - p_{wb}) \cdot (1 - p_c)$ |
| $m_6$ | Deleterious | Change to a different Deleterious Effect | $(1 - p_{wb}) \cdot p_c \cdot (1 - p_{cb})$ |
| $m_7$ | Deleterious | Change to Beneficial | $(1 - p_{wb}) \cdot p_c \cdot p_{cb}$ |

<sup>1</sup> $m_1$ are only defined at synonymous sites (1/3 of the total sites analyzed). $m_2$ to $m_7$ probabilities consider only non-synonymous sites (2/3 of the total sites analyzed). $p_{wb}$ is the fraction of mutations that are positively selected in the Wild population, $p_c$ is the fraction of mutations that change selection coefficient in the Domesticated population, and $p_{cb}$ is the fraction of those mutations that become beneficial in the Domesticated population (note in our simulations $p_{wb} = p_{cb}$ ). Note that in simulated scenarios $p_{cb} = p_{wb}$ .

### Nucleotide variability estimates

We have counted the number of polymorphic sites and estimated the Watterson variability estimate per nucleotide (Watterson 1975) for synonymous and non-synonymous sites for each of the 100 run simulations and for all the scenarios and populations. We have also calculated the ratio of synonymous versus non-synonymous polymorphic sites (*P_n_/P_s_*) as descriptive estimators of the observed variability at these sites (https://github.com/CastellanoED/domesticationDFE/blob/main/diversity_PnPs_slim.R).

### Distribution of fitness effects (DFE): Two complementary approaches

#### polyDFE: 1D-SFS and 1D-DFE

We use the polyDFEv2.0 framework (Tataru and Bataillon 2019) to estimate and compare the DFE across Wild-Domesticated population pairs by means of likelihood ratio tests (LRTs). We use the R function compareModels (from https://github.com/paula-tataru/polyDFE/blob/master/postprocessing.R) to compare pairs of models. The inference is performed only on the unfolded SFS data (divergence counts to the outgroup are not fitted), and unfolded SFS data are fitted using a DFE model comprising both deleterious (gamma distributed) and beneficial (exponentially distributed) mutations. The DFE of each Wild-Domesticated population pair is inferred using the 1D-SFS of each population. DFE is calculated assuming *S*=4*N*_e_*s*, in which *s* is the selective effect in the heterozygote, and *N*_e_ is the effective population size. Note that for comparative analysis *N*_e_ will be equivalent to *N*_a_. We used *S*/2 to contrast the simulated value with SLiM and with the inferred value in *dadi* (see next section). polyDFE assumes that new mutations in a genomic region arise as a Poisson process with an intensity that is proportional to the length of the region and the mutation rate per nucleotide (*μ*). We assume that *μ* remains constant across simulations (as it is the case). Both an ancestral SNP misidentification error (*ε*) and distortion parameters (*r_i_*) are estimated. However, we notice that the exclusion of *ε* does not affect the rest of estimated parameters because under the simulation conditions used here no sites are expected to be misidentified. The *r_i_* parameters are fitted independently for each frequency bin (from i = 1 to i = 19), and they are able to correct any distortion that affects equally the SFS of synonymous and non-synonymous variants (such as, in principle, demography or linked selection). Model averaging provides a way to obtain honest estimates that account for model uncertainty. To produce the model average estimates of the full DFE we weight each competing model according to their AIC following the equation 6.1 shown in the polyDFEv2 tutorial (“polyDFE/tutorial.pdf at master · paula-tataru/polyDFE”). We use the R function getAICweights (from https://github.com/paula-tataru/polyDFE/blob/master/postprocessing.R to do the model averaging) to obtain the AIC values.

#### dadi: 2D-SFS and 2D-DFE

*dadi* (Gutenkunst *et al*. 2009) is employed to infer the joint distribution of fitness effects (Jerison *et al*. 2014; Ragsdale *et al*. 2016; Huang *et al*. 2021) and the demographic history of all simulated population pairs. Following Huang *et al*. (2021), our model is that any mutation may have different selection coefficients *s*_w_ and *s*_d_ in the wild and domesticated populations, respectively. The joint DFE is the two-dimensional probability distribution quantifying the probability that a new mutation has selection coefficients *s_w_* and *s*_d_. In Huang *et al*. 2021, joint DFEs with only deleterious mutations were considered. Here we extend that model to consider joint DFEs that include mutations that are beneficial in one or both populations.

Our new model for the joint DFE between the two populations is a mixture of multiple components designed to mimic the selected mutation types in the simulations (Table 2; Figure 1D). The major exception is that beneficial mutations are modeled to have a single fixed selection coefficient, rather than arising from an exponential distribution. Let p_wb_ be the fraction of mutations that are positively selected in the Wild population, p_c_ be the fraction of mutations that change selection coefficient in the Domesticated population, and p_cb_ be the fraction of those mutations that become beneficial in the Domesticated population (note in our simulations p_wb_ = p_cb_). To model mutation types m_2_ and m_3_, a proportion p_wb_ · (1-p_c_) + p_wb_ · p_c_ · p_cb_ of mutations are assumed to have the same fixed positive selection coefficient in both populations. To model m_4_, a proportion p_wb_ · p_c_ · (1-p_cb_) is assumed to have a fixed positive selection coefficient in the Wild population and a gamma-distributed negative selection coefficient in the Domesticated population. To model m_5_, a proportion (1-p_wb_) · (1-p_c_) of mutations are assumed to have equal negative gamma-distributed selection coefficients in the two populations. To model m_6_, a proportion (1-p_wb_) · p_c_ · (1-p_cb_) is assumed to have independent gamma-distributed selection coefficients in the two populations. To model m_7_, a proportion (1-p_wb_) · pc · p_cb_ mutations is assumed to have a gamma-distributed negative selection coefficient in the Wild population and a fixed positive selection coefficient in the Domesticated population. All gamma distributions are assumed to have the same shape and scale. This model is implemented in *dadi* as the function **dadi.DFE.Vourlaki_mixture** (https://github.com/CastellanoED/domesticationDFE/blob/main/domestication_new_dadi_functions.py). Note in our simulations the marginal 1D-DFEs of Wild and Domesticated populations are exactly the same; the difference is that in Domesticated populations a given fraction of sites (some already polymorphic, some still monomorphic) can change their selection coefficient relative to the Wild population.

To infer the parameters of the joint DFE, we followed the procedure of Huang et. al (2021), but with this new DFE model. Briefly, assuming independence between mutations, the expected joint site frequency spectrum (SFS) for all mutations experiencing selection (here nonsynonymous sites) can be computed by integrating the joint SFS for each possible pair of population-size-scaled selection coefficients *S*_w_ and *S*_d_ over the joint DFE. Given that expected SFS, the composite likelihood of the nonsynonymous data can be computed by treating it as a Poisson Random Field, as in Gutenkunst et al. (2009). The parameters of the joint DFE model can then be inferred by maximizing that likelihood, using numerical optimization. For this study, we used the default in dadi, the BOBYQA optimization algorithm as implemented by the NLOpt library. For each selection coefficient pair (*S*_w_, *S*_d_), the expected SFS was calculated from a single integration of the partial differential equation (PDE) implemented by Gutenkunst et. al (2009) in the dadi software.

We integrate over our joint DFE model (Fig. 1D) by summing contributions for the discrete components of the DFE. The m_2_ + m_3_ component is simplest, being simply a weighting of the single SFS corresponding to the two positive selection coefficients assumed in the wild and domesticated populations. The m_4_ and m_7_ components are integrated over by holding *s*_w_ or *s*_d_ fixed and integrating over spectra calculated as the other selection coefficient is varied. The m_5_ component is integrated over by considering spectra in which *S*_w_ = *S*_d_. The m_4_, m_7_, and m_5_ components are thus one-dimensional integrations and employ the numerical methods developed in Kim et al. (2017). The m_6_ component is a two-dimensional integration over independent gamma distributions and is carried out as in Huang et al. (2021). This complex summation over spectra to calculate the expected SFS under the DFE is much less computationally expensive than calculating the spectra for each (*S*_w_, *S*_d_) pair, so those spectra are precomputed and cached.

For inference, a new, more general demographic model with branch-independent population size changes is first fit to the synonymous mutations from each simulation, and then the newly proposed joint DFE model is fit to the non-synonymous mutations. This model (Fig 1D) is implemented as a custom model using the dadi software and evaluated using the approach of Gutenkunst et al. (2009). The one subtlety is that an if statement is used to enable flexibility as to whether T1F or T2F is larger (https://github.com/CastellanoED/domesticationDFE/blob/main/domestication_new_dadi_functions.py, function name: “Domestication_flexible_demography”). The parameters of the demographic model (Figure 1B) are estimated by running 100 optimizations per inference unit. The 2D-SFS for selected sites are precomputed conditional on the demography for 104^2^ values of *S*_w_ and *S*_d_ *(S =*2*N*_a_s, a population scaled selection coefficient for the heterozygote where *N_a_* is the ancestral population size), 102 negative and 2 positives. For the negative part of the DFE, γ values were logarithmically equally spaced between −2000 and −10^−4^. The expected DFE for selected sites can then be computed as a weighted sum over these cached spectra (Kim *et al*. 2017). The DFE parameters shape, scale, *p*_wb_, *p*_c_, and *p*_cb_ are then estimated by maximizing the Poisson likelihood of the simulated data, with the non-synonymous rate of mutation influx fixed to twice that inferred for neutral sites in the demographic history fit. For the DFE inference, optimization is repeated until the best three results are within 0.5 log-likelihood units. Ancestral state misidentification is modelled, however in our simulations no sites are expected to be misidentified.

For the purpose of this work, *dadi* software is downloaded and installed according to the instructions provided at the following link: https://bitbucket.org/gutenkunstlab/dadi/src/master/. Since *dadi* operates as a module of Python, the Anaconda3 and Spyder (Python 3.7, Rossum and Drake 2009; Anaconda 2016; Raybaut 2009) versions are used in this study.

### Inference units, and confidence intervals in demographic and DFE parameters

To obtain the sampling variance of parameter estimates and approximate confidence intervals, we use a bootstrap approach. We resample with replacement 100 times 20 independent simulation runs or chromosomal “chunks” (from a pool of 100 “chunks”) and concatenate them. Hence, each concatenated unit (or inference unit) is made of 24 Mb of coding sequence (as comparison, the human genome contains ∼26 Mb of coding sequence). Uncertainties of DFE parameter inferences in polyDFE and *dadi* are calculated by this conventional bootstrapping, but in *dadi* we hold the demographic model fixed. In polyDFE the distortion introduced by demography (and linked selection) is not estimated but corrected with the *r_i_* parameters. Note that our procedure with *dadi* does not propagate uncertainty in demographic parameters through to the DFE parameters. To obtain the sampling variance of demographic parameter estimates with *dadi* we use the Godambe approach as described in Coffman *et al*. 2016. A final consideration on the factor of two differences across simulation and inference tools. We adjusted the population scaled selection coefficients to 2*N*_a_*s* in polyDFE, *dadi* and SLiM4 to enable a comparative study.

## RESULTS AND DISCUSSION

Studying the effect of domestication on the DFE of natural populations is particularly challenging, especially when available methods for inferring and comparing the DFE have not been evaluated using exactly the same dataset. In this study, we conduct simulations using different combinations of parameters relevant to the domestication process. A key distinction between the domestication demographic model used here and those commonly applied in speciation studies is the time scale since the split occurred. In our simulations, domesticated populations experience either large or small changes in the number and selective effects of loci under domestication, following a bottleneck period, with or without migration. Hereafter, we refer to these as the Wild and Domesticated populations.

This study focuses on the evolutionary process of domestication from the point of divergence to the present domesticated lineages. We do not account for the genetic improvement programs implemented in recent decades for some domesticated animals, which can significantly increase inbreeding levels (e.g., Makanjuola et al. 2020 estimated inbreeding levels as high as 40% in certain cattle breeds subjected to intense genomic selection). The simulated models in this work include strong selection and reductions in population size, both of which can moderately increase inbreeding levels in our simulations. However, Gilbert et al. (2022) reported that only very high selfing levels (>80%) severely affect DFE inference.

The Wild populations have a constant DFE and constant population size, but limited recombination across loci to mimic a realistic recombination landscape. Beneficial mutations arise at Wild populations following an exponential distribution, while deleterious mutations are drawn from a gamma distribution with shape 0.3 and mean S = −100 (where S = 2N_a_s, the selection coefficient *s* in the heterozygote is −1%, and N_a_ = 5,000 diploid individuals is the ancestral effective population size, see Material and Methods: Simulating the Domestication Process). As indicated in Materials and Methods section, all mutations, beneficial and deleterious, are co-dominant. The Domesticated population originates from the Wild population through a bottleneck and a concomitant change in selective effects at a fraction of non-synonymous sites (Figure 1; Table 1). The recombination and mutation landscapes are drawn from the same distribution in the Domesticated and Wild populations.

The change in selective effects affects both new mutations that arise within the Domesticated population and existing variants that existed before the domestication event. Put simply, not only can mutations that were deleterious (or beneficial) before the population split become beneficial (or deleterious) within the domesticated population, but even if the direction of the selective effect remains the same, the intensity of selection can change. Table 2 shows all the combinations of changes in selective effects between Wild and Domesticated populations. Our simulated scenarios aim to cover a variety of possible changes in the genetic architecture (number of loci) and the strength of selection (selection coefficients) of the trait/s under domestication. Three DFEs for beneficial mutations are assumed: (i) pervasive and nearly neutral, where a large fraction of new mutations (10%) are on average nearly neutral (*S*_b_ = 1), (ii) common and weak, where beneficial mutations are still fairly common (1%) but weakly selected (*S*_b_ = 10) and (iii) rare and strong, where very few mutations (0.1%) are strongly beneficial (*S*_b_ = 100). To better understand the role of selective sweeps on downstream inference, we also include simulations without a positive DFE. Depending on the scenario, a selective change occurs only at a small (0.05) or at a substantial proportion (0.25) of sites in the Domesticated population (Table 1, “*p*_c_” column). We leave eight scenarios as negative controls; the selection coefficients of new and standing variation in the Domesticated and Wild populations are exactly the same. Finally, demographic changes affect only the Domesticated population; the Wild population evolves under a constant population size. Two versions of the same demographic model (Figure 1A) are simulated: (i) one with migration, and (ii) another without migration. When there is migration, it only occurs from the Wild to the Domesticated population during the domestication bottleneck.

### Estimation of demographic parameters in Wild and Domesticated populations

In this study, we investigate the effects of natural selection—both broadly and in terms of how artificial selection alters the selective pressures acting on new and shared genetic variation— on the inference of demographic history and DFE during domestication. We do this using two commonly used inference tools (polyDFE and *dadi*) that assume free recombination across loci. Note that *dadi* first infers the demographic history and then infers the DFE assuming those inferred demographic parameters, whereas polyDFE operates independently of specific demographic histories and is designed to correct for distortions that affect both synonymous and non-synonymous site frequency spectra equally (Tataru and Bataillon 2019). Figure 1 A and B show the simulated joint demographic model and the joint demographic model used in the *dadi* inferences, respectively. We have increased the complexity of the inference model by introducing additional parameters, allowing it to account not only for “simulated” or true demographic changes, but also for more complex and unknown demographic histories and the potential influence of linked selection on synonymous SFS. The diagnostic plots can be found in Supplementary Figure 1; there is good agreement between the model fits and the data.

Our findings indicate that when positive selection is absent or relatively weak (S_b_ = 0, S_b_ = 1 or S_b_ = 10), the estimated onset of domestication tends to be approximately twice as old as the actual simulated starting point. Additionally, the inferred bottleneck appears slightly shallower but considerably longer than the simulated value (see Figure 2 and Supplementary Table 1 for the confidence intervals). This suggests that the influence of linked selection, likely driven primarily by background selection, has the effect of elongating the inferred timeline. Consequently, it makes the inferred domestication divergence and bottleneck appear more ancient and extended, respectively. For the Wild populations we always infer a larger population expansion than for the Domesticated populations, but without a bottleneck. This signal of a recent expansion in the Wild population is expected because when we consider how linked selection affects the SFS, there are more rare synonymous polymorphisms compared to what we would expect if there was free recombination under a constant population size (Charlesworth et al. 1993, 1995, Nielsen 2005, Zeng and Charleswoth 2011, Messer and Petrov 2013, Nicolaisen and Desai 2013, Ewing and Jensen, 2016). Remarkably, when positive selection is rare and strong (S_b_ = 100), the inferred temporal stretch becomes even more pronounced, and the inferred demographic history of both populations overlap extensively. The inferred domestication divergence shifts to approximately 50,000 years ago, whereas the actual simulated split occurred 10,000 years ago. Additionally, the inferred bottleneck appears significantly longer and less severe, while there is an inferred large population expansion in both Wild and Domesticated populations. Although in Figure 2 there appears to be a change in population size before the domestication split, only five scenarios (with IDs 3, 7, 15, 17 and 18) are statistically significant (Supplementary Table 1 and Supplementary Figure 2). Interestingly, we find the migration rate from Wild to Domesticated (*m*_w2d_) and from Domesticated to Wild (*m*_d2w_) are overestimated in most scenarios (Supplementary Table 1 and Supplementary Figure 3). We observe that neither migration nor an increase in *p*_c_ appears to significantly change the inferred demographic histories that we have just described.

**Figure 2.**
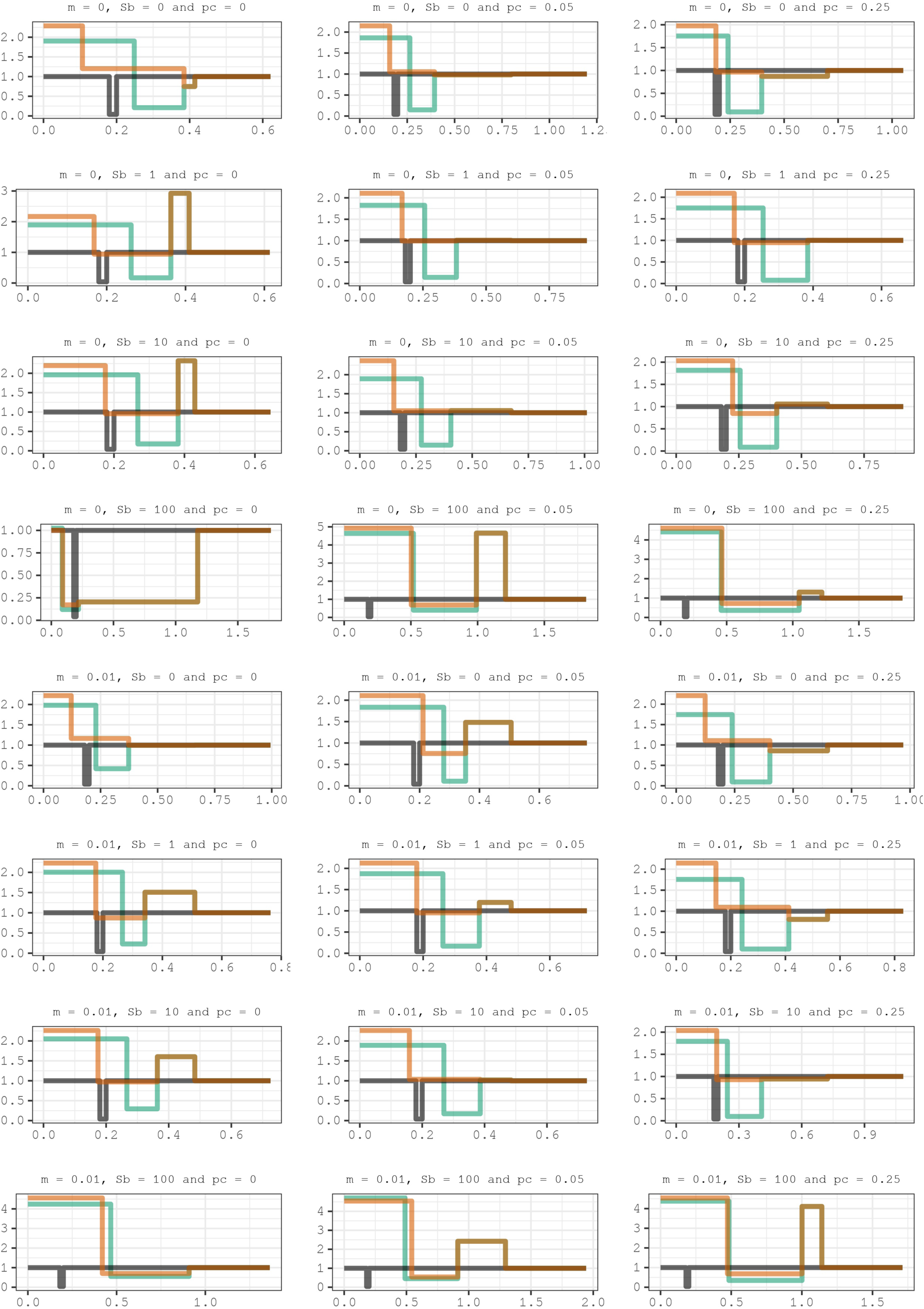
Lines showing the inferred demographic histories for the twenty-four simulated scenarios. In salmon-orange color is represented the Wild population and in turquoise-green color the Domesticated population. The dark grey line shows the true simulated demography in Domesticated populations. The true Wild population is not shown but it is a constant population size with relative *N_e_* = 1. The x-axis indicates the number of generations in relation to the ancestral population size *N_a_*, while the y-axis show the population size at each time in relation to *N_a_* (that is, *N_e_/N_a_*, where 1 means that *N_e_=N_a_*). The 95% confidence intervals calculated using the Godambe approximation can be found in Supplementary Table 1.

In summary, it was not possible to accurately determine the timing of the onset of domestication, the duration of the domestication bottleneck, or to distinguish between the presence and absence of migration between populations. We believe that these aspects are crucial for contextualizing the role of domestication in human history, and vice versa. Unfortunately, either the 2D-SFS or our “free recombination” modeling assumptions (or both) do not seem to be useful in this context.

Beyond domestication, the signal interference between selective and demographic processes has been widely studied. Linked selection significantly distorts the SFS, leading to biases in inferred demographic parameters. For example, Schrider et al. (2016) found that positive selection can mislead demographic inference, even inferring population size changes where none occurred, with selective sweeps as the primary cause. Gilbert et al. (2022) used forward simulations to report that large population expansions are inferred due to linked selection, particularly in regions of low recombination or high gene density. Finally, Johri et al. (2021) demonstrated biases due to background selection even after masking functional regions.

Together with these other findings, our work underscores the persistent difficulty of accurately inferring demographic histories in the presence of linked selection using population genomic data, even when using ancestral recombination graph based approaches (Marsh and Johri 2024).

Thus, the next question is to what extent can the nuisance *r_i_* parameters from polyDFE or this distorted inferred demography from *dadi* help to recover the simulated DFE parameters?

### Is it possible to detect domestication as an artificial change in the marginal full DFE between the two populations?

Next, we investigate whether polyDFE captures differences in the marginal (or 1D) full DFE of Domesticated and Wild populations across the twenty-four domestication scenarios (Table 1). We run five nested models (Table 3) and compare them using likelihood ratio tests (LRTs) (Supplementary Table 2). It is important to note that in all our simulations, the marginal full DFE for new mutations in both Domesticated and Wild populations is the same within a given scenario (as detailed in Table 1). This means that the selection coefficients for sites, whether they are monomorphic or polymorphic, are drawn from the same full DFE. In simpler terms, the proportion of new mutations that are advantageous or detrimental is identical for both Domesticated and Wild populations within a given scenario.

**Table 3.**
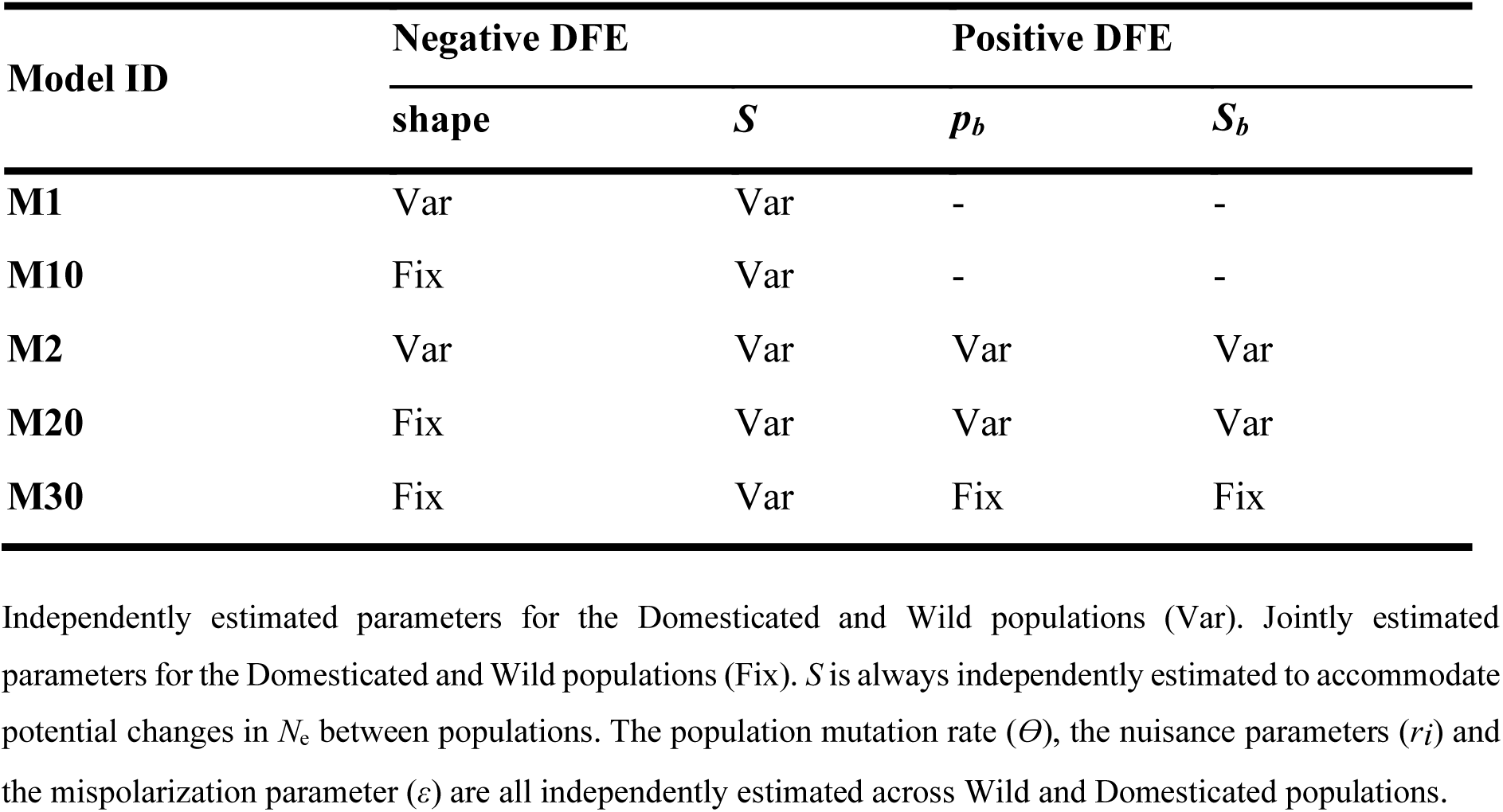
List of nested polyDFE models and (co)estimated parameters.

LRTs between different nested models allow us to address important questions about the DFE, without assuming any prior knowledge of our datasets. First, we assess whether the inferred shape of the negative DFE is similar in both populations while also examining if the estimation of the shape parameter is influenced by the presence of advantageous mutations. When comparing models that do not consider beneficial mutations (models M1 versus M10 in the second column of Supplementary Table 2), the model with a distinct shape for Domesticated and Wild populations is accepted only in two, rather unrelated, scenarios (scenarios 7 and 11). This indicates that an artificial alteration in the shape of the deleterious DFE between Domesticated and Wild populations can be inferred. Fortunately, when comparing models that take into account beneficial mutations (models M2 vs M20, third column in Supplementary Table 2), all scenarios show a shared shape of the deleterious DFE, which is expected based on the simulation parameters. These findings suggest that disregarding beneficial mutations can cause an artificial change in the inferred shape of the marginal deleterious DFE between populations, as noted previously by Tataru et al. in 2017. Second, when we contrast models with and without considering the positive DFE (that is, testing the nested models M1 vs M2 and M10 vs M20), yields statistically significant results in all scenarios (see Supplementary Table 2, fourth and fifth columns). Hence, polyDFE appears to detect beneficial mutations, regardless of the true presence and strength of positive selection. Third, we investigate whether Domesticated and Wild populations could exhibit an artificial change in the beneficial DFEs as a consequence of domestication. When comparing the M20 and M30 models (refer to the last column in Supplementary Table 2), polyDFE invokes changes in the positive DFE between populations in most scenarios without migration (with IDs 1, 2, 5, 7, 8, 11 and 12). Below we characterize this putative change in the marginal DFEs between populations.

### Estimation of DFE parameters in Wild and Domesticated populations

Under the polyDFE framework, we begin by extracting the Akaike Information Criterion (AIC) from every model (Table 3) and then computing the AIC-weighted parameters for all models (Tataru and Bataillon 2019; Castellano *et al*. 2019). This approach is used because the true model generating real data in both Wild and Domesticated populations is unknown. Instead, under *dadi*’s framework, we adopt an alternative methodology that utilizes very general, parameter rich and versatile joint demographic and DFE models to fit the 2D-SFS. The diagnostic plots of the new joint DFE model is shown in Supplementary Figure 1, again there is good agreement between the model fits and the data.

Inferred parameters related to the deleterious DFE: Supplementary Figure 4 and 5 depicts the distribution of parameters related to the deleterious DFE that are estimated by performing bootstrap analysis using polyDFE and *dadi.* We observe that both tools have a tendency to marginally overestimate the shape parameter of the gamma distribution employed to model the deleterious DFE (Supplementary Figure 4). The overestimation is particularly significant in polyDFE, when positive selection is strong. In such scenarios, *dadi*’s shape estimation is sometimes rather noisy. Regarding the mean of the deleterious DFE (*s*) (Supplementary Figure 5), we observe that the inferred mean values across bootstrap replicates vary by up to 20% higher or lower, depending on the population, scenario, and inference tool. The largest misinference occurs when positive selection is strong and *dadi* is used and in the Domesticated population when polyDFE is used.

Inferred parameters related to the beneficial DFE: The distribution of parameters associated with the beneficial DFE, estimated by bootstrap analysis using polyDFE and *dadi*, is shown in Supplementary Figure 6 and Supplementary Table 3 (only *dadi*). Depending on the scenario, we simulate an average increase in relative fitness (*s*_b_) of 0.010, 0.001, and 0.0001. Positive selection’s strength is usually substantially underestimated by polyDFE and *dadi*, but only polyDFE consistently overestimates the proportion of new advantageous mutations (*p*_b_), regardless of the true simulated value. Given the distribution of inferred values of *p_b_* and *s_b_*, a peak of effectively neutral advantageous mutations is being measured by polyDFE. The overall excess of effectively neutral advantageous mutations measured by polyDFE is generally balanced by the defect of effectively neutral deleterious mutations. Consequently, polyDFE seems to have limited power in identifying effectively beneficial mutations on the 1D-SFS (under these simulation conditions). More importantly, the apparent spurious difference in the marginal full DFE between populations detected by polyDFE disappears when the full DFE is discretized. We conclude that if a significant change is detected in the discretized marginal full DFE, it must be considered valid.

It is noteworthy that both polyDFE and *dadi* tools typically produce comparable and reasonably accurate discretized deleterious DFEs (Figure 3), despite polyDFE’s tendency to infer a peak of effectively neutral beneficial mutations. This suggests that, regardless of the inference method used, the estimation of the “effective” discretized deleterious DFE remains robust to demographic and selective changes, as well as the pervasive effects of linked selection. In contrast, recent studies indicate that in highly selfing species, the deleterious DFE is often misestimated due to the influence of linked selection (Gilbert et al. 2022), particularly strong Hill-Robertson interference (Daigle and Johri 2024). These findings highlight that the accuracy of inferring the deleterious DFE is not universal but instead depends on factors such as the degree of selfing and inbreeding.

**Figure 3.**
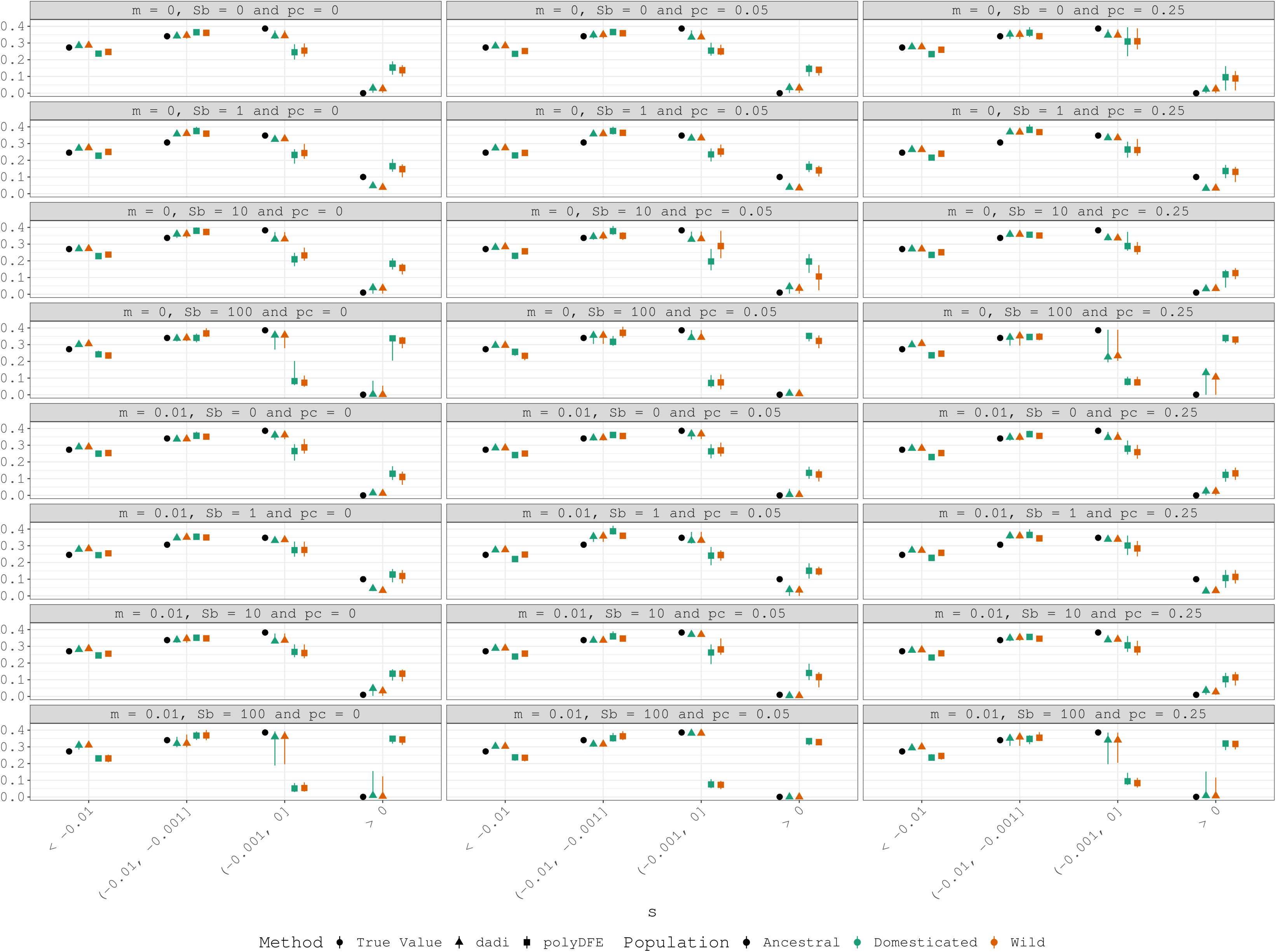
Sampling distributions of estimated discretized full DFE obtained using 100 bootstrap replicates.

Thus, we conclude that both tools struggle to infer the positive DFE and tend to be overconservative and identify weaker positive selection than what has been simulated. We suspect this arises from linkage between beneficial and synonymous mutations, which may lead to an excess of high-frequency synonymous mutations and an overcorrection of the excess non-synonymous polymorphisms at high frequency, either through polyDFE’s *r*_i_ parameters or *dadi*’s inferred demographic history. Notably, these findings are consistent with what was already pointed out by Tataru et al. (2017) and Booker et al (2020) using a single population. They draw attention to the challenge of inferring parameters of positive selection when counting for weakly and strongly selected mutations. Indeed, Booker et al. (2020) emphasize that in the case of rare and strong positive selection, the SFS can be very noisy, with linked sites playing an important role, making it difficult to infer the positive DFE.

### Estimation of the fraction of mutations with divergent selective effects (*p_c_*) between Domesticated and Wild populations

One of the main goals of this study is to determine the proportion of new and standing non-synonymous mutations with differing selection coefficients in Wild and Domesticated populations. The usage of our new joint DFE model is not limited to the current study. Our new model, created by mixing multiple distributions to mimic mutation types in our simulations (Table 2; Figure 1C-D), is suitable for usage in any recently diverged populations.

Figure 4 displays the distribution of the inferred *p_c_* for three different positive DFEs, along with simulated *p*_c_ values. When positive selection is not strong, it becomes apparent that scenarios with a significant fraction of mutations with dissimilar selective effects (p_c_ = 0.25) can readily be differentiated from those where a small (p_c_ = 0.05) or nonexistent (*p_c_* = 0) number of sites alter their selection coefficient. However, differentiating our negative control from a positive control proves difficult when only 0.05 of the sites show a difference in their selection coefficients. Notably, we overestimate *p_c_* significantly in cases of strong positive selection, indicating that classic hard selective sweeps may mimic divergent selection in a substantial amount of non-synonymous mutations. We observe no major impact of migration on the inferred *p*_c_ values across scenarios.

**Figure 4.**
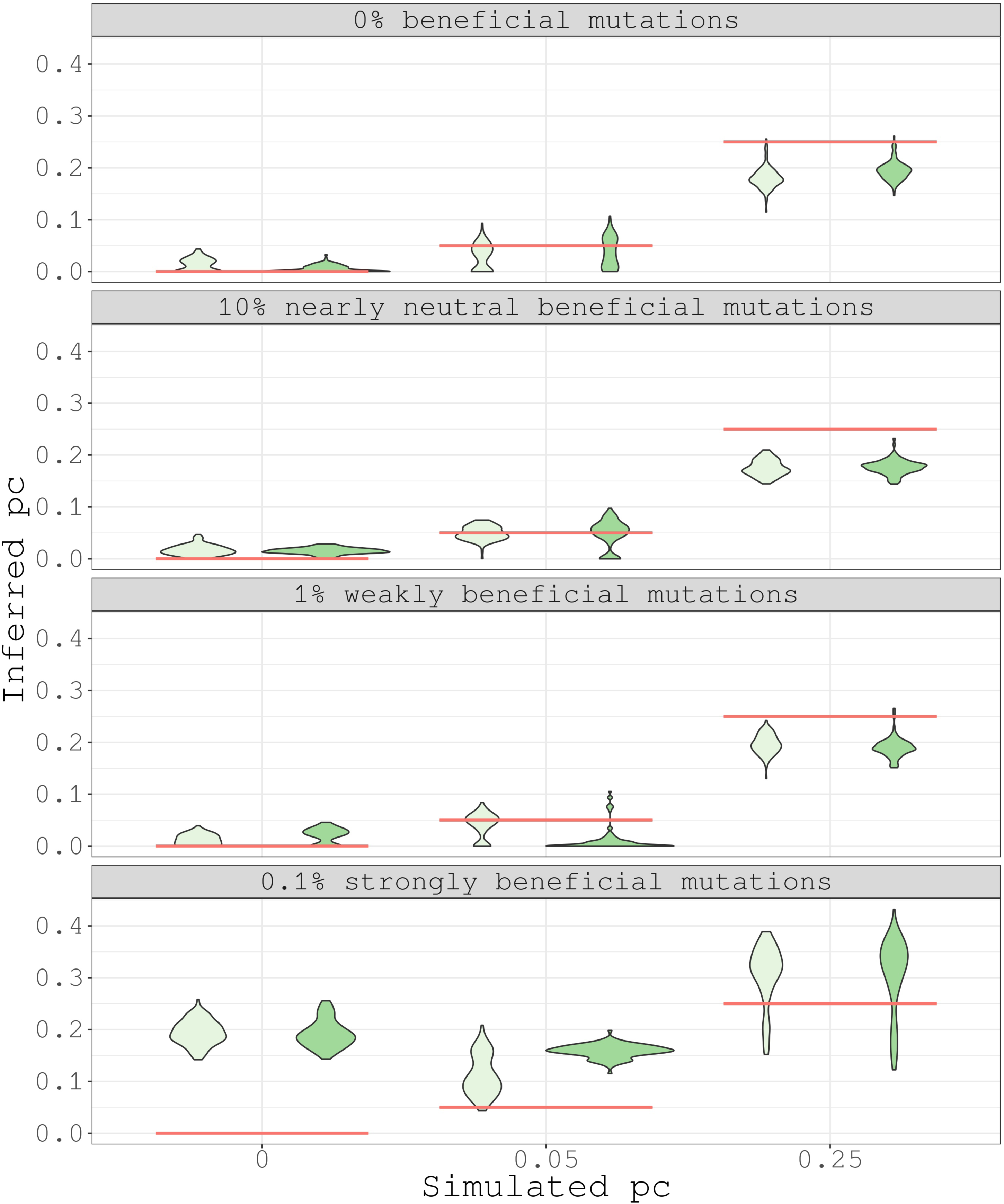
Sampling distributions of inferred *p_c_* (dadi) are obtained using 100 bootstrap replicates. In light green are shown the scenarios without migration and in dark green the scenarios with migration.

The overestimation of *p_c_* when positive selection is strong is not surprising, since non-synonymous mutations with stable selection coefficients between populations may be in close recombinational proximity and hitchhike with strongly beneficial mutations that are population-specific. This will exacerbate the apparent fraction of mutations with divergent selective effects. In contrast, if positive selection is weaker, recombination will be able to disentangle beneficial mutations from the rest of mutation types and simplify our estimation of *p_c_*. One way to ameliorate this problem would be to remove genomic windows with evidence of recent population-specific, complete or partial, selective sweeps and rerun our inference pipeline. For example, these could be regions with low neutral genetic diversity. However, we find this heuristic solution might be difficult to implement in practice.

Supplementary Figure 7 shows the observed level of neutral genetic diversity (measured using Watterson’s theta (Watterson 1975) and synonymous sites, *θ_s_*) and the selective constraint (i.e., the ratio of non-synonymous polymorphisms to synonymous polymorphisms per site, *P_n_/P_s_*) for each independent simulation run. Note the large decrease in the observed *θ_s_*, driven entirely by linked selection in Wild populations, relative to the expected level of neutral genetic diversity (expected *θ_s_* = 0.005 under free recombination). Particularly important is the reduction in the average *θ_s_* across independent simulation runs in Wild populations when positive selection is rare and strong (*θ_s_* is ∼20% of the expected value), whereas when positive selection is weaker or absent the observed level of genetic diversity is ∼40% of the expected value. In strong positive selection scenarios, there may be no heuristic correction or genomic region that escapes *genetic draft* (Gillespie 2000), and our current definition and interpretation of *p*_c_ would be misleading. We also observe that when positive selection is strong, genetic diversity and *P_n_/P_s_* are significantly further reduced in both Domesticated and Wild populations, causing the two distributions to largely overlap. As described above for the reconstruction of demographic history, when selective sweeps are strong, the recovered demographic history also tends to overlap between Wild and Domesticated populations. The overlap of demographic histories, neutral genetic diversity and *P_n_/P_s_* distributions could be used as a caution signal and as an indicator of strong positive selection and widespread genetic draft. Finally, migration appears to cause a minor reduction in *P_n_/P_s_* and increase genetic diversity within Domesticated populations. Thus, migration acts slightly diminishing the *P_n_/P_s_* discrepancy between Wild and Domesticated populations.

### Implications for empirical analysis of populations

A scenario involving divergent populations, with one undergoing a bottleneck and a shift in the selection regime, may also be relevant in other contexts beyond domestication, such as invasive species, island colonization or recent parapatric and allopatric speciation events. In this work, we observed that selective effects affect the inference of demographic parameters by linked selection, but to different extents depending on the DFE. Background selection contributes to the misinference of domestication divergence time and the duration of the bottleneck, making them appear more ancient and extended than in our simulations. When strong selective sweeps are combined with background selection, the inferred temporal stretch becomes even more pronounced, and the inferred demographic history of both populations overlaps extensively. These demographic distortions in the inference must be considered when interpreting real data using these methods or any other methods that make similar assumptions. Nevertheless, under the assumptions used in this work, we believe that the discretized deleterious DFE is estimated with reasonable accuracy. This suggests that methods designed to infer the entire DFE could be applied first, followed by the estimation of demographic parameters using this information. Interestingly, Johri et al. (2021), using a different approach based on a single population and considering four classes of deleterious mutations, found that while DFE classes were accurately estimated, demographic parameters were not. They proposed a method to jointly infer both demography and deleterious mutations using an ABC framework. Although computationally intensive, this approach may help address some of the inference challenges highlighted in this work.

Another point of interest for empirical geneticists is the development of a new method to jointly infer the DFE between wild and domesticated and their differences in the positive part of the distribution. The 2D *dadi* extension algorithm allows to infer differences in *p*_wb_ (the fraction of mutations that are positively selected in the wild population), *p*_c_ (the fraction of mutations that change the coefficient of selection in the domesticated population), *p*_cb_ (the fraction of those mutations that become beneficial in the domesticated population).

## CONCLUSIONS

In summary, our use of forward-in-time simulations has provided valuable insights into the inference of complex genetic demographic history and distribution of fitness effects (DFE) for both new and standing amino acid mutations in the context of domestication. Through a comparative analysis of two methods, polyDFE and *dadi*, and the new implementation of a full 2D-SFS full inference of DFE, we have uncovered the impact of linked selection on the reconstructed demographic history of both wild and domesticated populations. Despite biases in the timelines of domestication events and bottleneck characteristics, the estimation of discretized deleterious DFE remains remarkably reliable, demonstrating the robustness of these analytical approaches in the studied conditions. In particular, the underestimation of effectively beneficial mutations in the DFE highlights the influence of linkage between beneficial and neutral mutations, which requires further consideration in model design and interpretation. In addition, our results shed light on distinguishing scenarios of divergent selective effects between populations under weak and strong positive selection, providing a nuanced understanding of the interplay of evolutionary forces. Nevertheless, we must approach the results of this work with caution, as the simulated demographic and selective patterns are based on specific models/idealizations that may not fully capture the complexities of domestication. On the other hand, as we navigate the complex landscape of domestication, these methodological approaches contribute significantly to unraveling the evolutionary dynamics and adaptive processes that shape the genomes of domesticated organisms, and provide a foundation for future research in this critical area of study.

## Supplementary Information

We have added the Diagnostic plots 2D-SFS for synonymous and Nonsynonymous positions obtained with *dadi* for all the scenarios analyzed, plus seven additional Figures and three additional Tables in the Supplementary Material. This Supplementary Information is available at Zenodo: https://zenodo.org/records/14277802, as well as the scripts used in the analyses of this work.

## LITERATURE CITED

Ahmad H. I., M. J. Ahmad, F. Jabbir, S. Ahmar, N. Ahmad, et al., 2020 The Domestication Makeup: Evolution, Survival, and Challenges. Front. Ecol. Evol. 8.

Amills M., H. J. W. C. Megens, A. Manunza, S. E. Ramos-Onsins, and M. Groenen, 2017 A genomic perspective about wild boar demography and evolution, pp. 376–387 in Ecology, Conservation and Management of Wild Pigs and Peccaries, Cambridge University Press.

Anaconda Software Distribution. Computer software. Vers. 2-2.4.0. Anaconda, Nov. 2016. Web.

Andersson L., 2012 Genetics of Animal Domestication, pp. 260–274 in Biodiversity in Agriculture: Domestication, Evolution, and Sustainability, edited by Damania A. B., Qualset C. O., McGuire P. E., Gepts P., Bettinger R. L., et al. Cambridge University Press, Cambridge.

Arnoux S., C. Fraïsse, C. Sauvage. Genomic inference of complex domestication histories in three Solanaceae species. J Evol Biol. 2021 Feb;34(2):270–283. 10.1111/jeb.13723. Epub 2020 Nov 10. PMID: 33107098.

Avni R., M. Nave, O. Barad, K. Baruch, S. O. Twardziok, et al., 2017 Wild emmer genome architecture and diversity elucidate wheat evolution and domestication. Science 357: 93–97. 10.1126/science.aan0032

Barton H. J., and K. Zeng, 2018 New Methods for Inferring the Distribution of Fitness Effects for INDELs and SNPs. Mol. Biol. Evol. 35: 1536–1546. 10.1093/molbev/msy054

Beissinger T. M., L. Wang, K. Crosby, A. Durvasula., M. B. Hufford, J. Ross-Ibarra. Recent demography drives changes in linked selection across the maize genome. Nat Plants. 2016 Jun 13;2:16084. 10.1038/nplants.2016.84. PMID: 27294617.

Booker T.R. 2020 Inferring Parameters of the Distribution of Fitness Effects of New Mutations When Beneficial Mutations Are Strongly Advantageous and Rare. G3 (Bethesda). 10(7):2317–2326. 10.1534/g3.120.401052.

Boyko A. R., R. H. Boyko, C. M. Boyko, H. G. Parker, M. Castelhano, et al., 2009 Complex population structure in African village dogs and its implications for inferring dog domestication history. Proc. Natl. Acad. Sci. U. S. A. 106: 13903–13908. 10.1073/pnas.0902129106

Caicedo A. L., S. H. Williamson, R. D. Hernandez, A. Boyko, A. Fledel-Alon, T. L. York, N. R. Polato, K. M. Olsen, R. Nielsen, S. R. McCouch, C. D. Bustamante, M. D. Purugganan. Genome-wide patterns of nucleotide polymorphism in domesticated rice. PLoS Genet. 2007 Sep;3(9):1745–56. 10.1371/journal.pgen.0030163. Epub 2007 Aug 6.

Carneiro M., Piorno V., Rubin C.J., Alves J.M., Ferrand N., Alves P.C., Andersson L. 2015 Candidate genes underlying heritable differences in reproductive seasonality between wild and domestic rabbits. Anim Genet. 418–25. 10.1111/age.12299

Castellano D., M. C. Macià, P. Tataru, T. Bataillon, and K. Munch, 2019 Comparison of the Full Distribution of Fitness Effects of New Amino Acid Mutations Across Great Apes. Genetics 213: 953–966. 10.1534/genetics.119.302494

Charlesworth, B., M. Morgan, and D. Charlesworth, 1993 The effect of deleterious mutations on neutral molecular variation. Genetics 134: 1289. 10.1093/genetics/134.4.1289

Charlesworth D., B. Charlesworth., M. T. Morgan. 1995 The pattern of neutral molecular variation under the background selection model. Genetics. 141(4):1619–32. 10.1093/genetics/141.4.1619.

Coffman A. J., P. H. Hsieh, S. Gravel, and R. N. Gutenkunst, 2016 Computationally Efficient Composite Likelihood Statistics for Demographic Inference. Mol. Biol. Evol. 33: 591–593. 10.1093/molbev/msv255

Comeron J.M. Background selection as null hypothesis in population genomics: insights and challenges from Drosophila studies. Philos Trans R Soc Lond B Biol Sci. 2017 Dec 19;372(1736):20160471. 10.1098/rstb.2016.0471. PMID: 29109230; PMCID: PMC5698629.

Courtier-Orgogozo V., and A. Martin, 2020 The coding loci of evolution and domestication: current knowledge and implications for bio-inspired genome editing, (M. H. Dickinson, 27 L. B. Vosshall, and J. A. T. Dow, Eds.). J. Exp. Biol. 223: jeb208934. 10.1242/jeb.208934

Crow J. F., and M. Kimura, 1979 Efficiency of truncation selection*. Proc. Natl. Acad. Sci. U. S. A. 76: 396–399.

Dayan T., 1994 Early Domesticated Dogs of the Near East. J. Archaeol. Sci. 21: 633–640. 10.1006/jasc.1994.1062

Daigle A., P. Johri. 2024 Hill-Robertson interference may bias the inference of fitness effects of new mutations in highly selfing species. bioRxiv [Preprint].12:2024.02.06.579142. 10.1101/2024.02.06.579142.

Ewing G. B., J. D. Jensen. 2016 The consequences of not accounting for background selection in demographic inference. Mol Ecol. 25(1):135–41. 10.1111/mec.13390.

Frantz L. A. F., J. G. Schraiber, O. Madsen, H.-J. Megens, A. Cagan, et al., 2015 Evidence of long-term gene flow and selection during domestication from analyses of Eurasian wild and domestic pig genomes. Nat. Genet. 47: 1141–1148. 10.1038/ng.3394

Galtier N. Adaptive Protein Evolution in Animals and the Effective Population Size Hypothesis. PLoS Genet. 2016 Jan 11;12(1):e1005774. 10.1371/journal.pgen.1005774. PMID: 26752180; PMCID: PMC4709115.

Gilbert K.J., S. Zdraljevic, D. E. Cook, A. D. Cutter, E. C. Andersen, C. F. Baer. The distribution of mutational effects on fitness in Caenorhabditis elegans inferred from standing genetic variation. Genetics. 2022 Jan 4;220(1):iyab166. 10.1093/genetics/iyab166.

Gillespie J. H., 2000 Genetic drift in an infinite population. The pseudohitchhiking model. Genetics 155: 909–919. 10.1093/genetics/155.2.909

Granleese T., S. A. Clark, and J. H. J. van der Werf, 2019 Genotyping strategies of selection candidates in livestock breeding programmes. J. Anim. Breed. Genet. 136: 91–101. 10.1111/jbg.12381

Groenen M. A. M., A. L. Archibald, H. Uenishi, C. K. Tuggle, Y. Takeuchi, et al., 2012 Analyses of pig genomes provide insight into porcine demography and evolution. Nature 491: 393–398. 10.1038/nature11622

Gutenkunst R. N., R. D. Hernandez, S. H. Williamson, and C. D. Bustamante, 2009 Inferring the Joint Demographic History of Multiple Populations from Multidimensional SNP Frequency Data. PLOS Genet. 5: e1000695. 10.1371/journal.pgen.1000695

Haller B. C., and P. W. Messer, 2023 SLiM 4: Multispecies Eco-Evolutionary Modeling. Am. Nat. 201: E127–E139. 10.1086/723601

Huang X., A. L. Fortier, A. J. Coffman, T. J. Struck, M. N. Irby, et al., 2021 Inferring Genome-Wide Correlations of Mutation Fitness Effects between Populations. Mol. Biol. Evol. 38: 4588–4602. 10.1093/molbev/msab162

Huang X., N. Kurata, X. Wei, Z.-X. Wang, A. Wang, et al., 2012 A map of rice genome variation reveals the origin of cultivated rice. Nature 490: 497–501. 10.1038/nature11532

Jain K., and W. Stephan, 2017a Rapid Adaptation of a Polygenic Trait After a Sudden Environmental Shift. Genetics 206: 389–406. 10.1534/genetics.116.196972

Jain K., and W. Stephan, 2017b Modes of Rapid Polygenic Adaptation. Mol. Biol. Evol. 34: 3169–3175. 10.1093/molbev/msx240

Jerison E. R., S. Kryazhimskiy, and M. M. Desai, 2014 Pleiotropic consequences of adaptation across gradations of environmental stress in budding yeast. arXiv:1409.7839, 10.48550/arXiv.1409.7839

Johri P., K. Riall, H. Becher, L. Excoffier, B. Charlesworth, J. D. Jensen, 2021 The Impact of Purifying and Background Selection on the Inference of Population History: Problems and Prospects. Mol Biol Evol. 38(7):2986–3003. 10.1093/molbev/msab050.

Keightley P. D., and A. Eyre-Walker, 2007 Joint inference of the distribution of fitness effects of deleterious mutations and population demography based on nucleotide polymorphism frequencies. Genetics 177: 2251–2261. 10.1534/genetics.107.080663

Kim B. Y., C. D. Huber, and K. E. Lohmueller, 2017 Inference of the Distribution of Selection Coefficients for New Nonsynonymous Mutations Using Large Samples. Genetics 206: 345–361. 10.1534/genetics.116.197145

Kondrashov A. S., 1988 Deleterious mutations and the evolution of sexual reproduction. Nature 336: 435–440. 10.1038/336435a0

Larson G., and J. Burger, 2013 A population genetics view of animal domestication. Trends Genet. TIG 29: 197–205. 10.1016/j.tig.2013.01.003

Larson G., D. R. Piperno, R. G. Allaby, M. D. Purugganan, L. Andersson, et al., 2014 Current perspectives and the future of domestication studies. Proc. Natl. Acad. Sci. 111: 6139–6146. 10.1073/pnas.1323964111

Leno-Colorado J., S. Guirao-Rico, M. Pérez-Enciso, and S. E. Ramos-Onsins, 2022 Pervasive selection pressure in wild and domestic pigs. bioRxiv 2020.09.09.289439; 10.1101/2020.09.09.289439

Li N., R. Xu, P. Duan, and Y. Li, 2018 Control of grain size in rice. Plant Reprod. 31: 237–251. 10.1007/s00497-018-0333-6

Li X, Yang J, Shen M, Xie XL, Liu GJ, Xu YX, et al. 2020. Whole-genome resequencing of wild and domestic sheep identifies genes associated with morphological and agronomic traits. Nat Commun. https://www.doi.org/10.1038/s41467-020-16485-1.

Librado P., N. Khan, A. Fages, M. A. Kusliy, T. Suchan, et al., 2021 The origins and spread of domestic horses from the Western Eurasian steppes. Nature 598: 634–640. 10.1038/s41586-021-04018-9

Lozano R., E. Gazave, J. P. R. Dos Santos, M. G. Stetter, R. Valluru, N. Bandillo, S. B. Fernandes, P. J. Brown, N. Shakoor, T. C. Mockler, E. A. Cooper, M. Taylor Perkins, E. S. Buckler, J. Ross-Ibarra, M. A. Gore. Comparative evolutionary genetics of deleterious load in sorghum and maize. Nat Plants. 2021 Jan;7(1):17–24. 10.1038/s41477-020-00834-5. Epub 2021 Jan 15. PMID: 33452486.

Lu J., T. Tang, H. Tang, J. Huang, S. Shi, et al., 2006 The accumulation of deleterious mutations in rice genomes: a hypothesis on the cost of domestication. Trends Genet. 22: 126–131. 10.1016/j.tig.2006.01.004

Makanjuola B. O., F. Miglior, E. A. Abdalla, C. Maltecca, F. S. Schenkel, C. F. Baes. Effect of genomic selection on rate of inbreeding and coancestry and effective population size of Holstein and Jersey cattle populations. J Dairy Sci. 2020 Jun;103(6):5183–5199. 10.3168/jds.2019-18013.

Marsden C. D., D. Ortega-Del Vecchyo, D. P. O’Brien, J. F. Taylor, O. Ramirez, et al., 2016 Bottlenecks and selective sweeps during domestication have increased deleterious genetic variation in dogs. Proc. Natl. Acad. Sci. 113: 152–157. 10.1073/pnas.1512501113

Marsh J.I., P. Johri. Biases in ARG-Based Inference of Historical Population Size in Populations Experiencing Selection. Mol Biol Evol. 2024 Jul 3;41(7):msae118. doi: 10.1093/molbev/msae118.

Messer P. W., D. A. Petrov. 2013 Frequent adaptation and the McDonald-Kreitman test. Proc Natl Acad Sci U S A. 110(21):8615–20. 10.1073/pnas.1220835110

Morell Miranda P., A. E. R. Soares, T. Günther. Demographic reconstruction of the Western sheep expansion from whole-genome sequences. G3 (Bethesda). 2023 Nov 1;13(11):jkad199. 10.1093/g3journal/jkad199. PMID: 37675574.

Moyers, Morrell, and McKay, 2018 Genetic Costs of Domestication and Improvement. J. Hered. 109. 10.1093/jhered/esx069

Murray C., E. Huerta-Sanchez, F. Casey, and D. G. Bradley, 2010 Cattle demographic history modelled from autosomal sequence variation. Philos. Trans. R. Soc. B Biol. Sci. 365: 2531–2539. 10.1098/rstb.2010.0103

Murray C, E. Huerta-Sanchez, F. Casey, D. G. Bradley. Cattle demographic history modelled from autosomal sequence variation. Philos Trans R Soc Lond B Biol Sci. 2010 Aug 27;365(1552):2531–9. 10.1098/rstb.2010.0103. PMID: 20643743; PMCID: PMC2935105.

Nicolaisen L.E., M. M. Desai. 2013 Distortions in genealogies due to purifying selection and recombination. Genetics. 195(1):221–30. 10.1534/genetics.113.152983

Nielsen R., 2005 Molecular signatures of natural selection. Annu. Rev. Genet. 39: 197–218. 10.1146/annurev.genet.39.073003.112420

Ohta T., 1989 The mutational load of a multigene family with uniform members. Genet. Res. 53: 141–145. 10.1017/s0016672300028020

Pacifici M., L. Santini, M. D. Marco, D. Baisero, L. Francucci, et al., 2013 Generation length for mammals. Nat. Conserv. 5: 89–94. 10.3897/natureconservation.5.5734

Purugganan M. D., and D. Q. Fuller, 2009 The nature of selection during plant domestication. Nature 457: 843–848. 10.1038/nature07895

Qanbari S., C.J. Rubin, K. Maqbool, S. Weigend, A. Weigend, J. Geibel et al. 2019 Genetics of adaptation in modern chicken. PLoS Genet. 15(4):e1007989. 10.1371/journal.pgen.1007989

Ragsdale A. P., A. J. Coffman, P. Hsieh, T. J. Struck, and R. N. Gutenkunst, 2016 Triallelic Population Genomics for Inferring Correlated Fitness Effects of Same Site Nonsynonymous Mutations. Genetics 203: 513–523. 10.1534/genetics.115.184812

Raybaut, P. (2009). Spyder-documentation. Available Online at: Pythonhosted. Org.

Redding R. W., 2015 The Pig and the Chicken in the Middle East: Modeling Human Subsistence Behavior in the Archaeological Record Using Historical and Animal Husbandry Data. J. Archaeol. Res. 23: 325–368. 10.1007/s10814-015-9083-2

Rossum G. V., and F. L. Drake, 2009 Python 3 Reference Manual: (Python Documentation Manual Part 2). CreateSpace Independent Publishing Platform.

Rubin C.-J., M. C. Zody, J. Eriksson, J. R. S. Meadows, E. Sherwood, et al., 2010 Whole-genome resequencing reveals loci under selection during chicken domestication. Nature 464: 587–591. 10.1038/nature08832

Schrider D.R., A.G Shanku., A.D Kern. 2016 Effects of Linked Selective Sweeps on Demographic Inference and Model Selection. Genetics. 204(3):1207–1223. 10.1534/genetics.116.190223.

Shomura A., T. Izawa, K. Ebana, T. Ebitani, H. Kanegae, et al., 2008 Deletion in a gene associated with grain size increased yields during rice domestication. Nat. Genet. 40: 1023–1028. 10.1038/ng.169

Stephan W., and S. John, 2020 Polygenic Adaptation in a Population of Finite Size. Entropy 22: 907. 10.3390/e22080907

Tataru P., and T. Bataillon, 2019 polyDFEv2.0: testing for invariance of the distribution of fitness effects within and across species. Bioinforma. Oxf. Engl. 35: 2868–2869. 10.1093/bioinformatics/bty1060

Tataru P., M. Mollion, S. Glémin, and T. Bataillon, 2017 Inference of Distribution of Fitness Effects and Proportion of Adaptive Substitutions from Polymorphism Data. Genetics 207: 1103–1119. 10.1534/genetics.117.300323

Tataru P. 2018. polyDFE/tutorial.pdf at master. https://github.com/paula-tataru/polyDFE

Todd E. T., L. Tonasso-Calvière, L. Chauvey, S. Schiavinato, A. Fages, et al., 2022 The genomic history and global expansion of domestic donkeys. Science 377: 1172–1180. 10.1126/science.abo3503

Torres R, M. G. Stetter, R.D. Hernandez, J. Ross-Ibarra. The Temporal Dynamics of Background Selection in Nonequilibrium Populations. Genetics. 2020 Apr;214(4):1019–1030. 10.1534/genetics.119.302892. Epub 2020 Feb 18. PMID: 32071195; PMCID: PMC7153942.

Van Laere A.-S., M. Nguyen, M. Braunschweig, C. Nezer, C. Collette, et al., 2003 A regulatory mutation in IGF2 causes a major QTL effect on muscle growth in the pig. Nature 425: 832–836. 10.1038/nature02064

Watterson G. 1975 On the number of segregating sites in genetical models without recombination. Theoretical Population Biology, 7, 256. 10.1016/0040-5809(75)90020-9.

Wright S. I., I. V. Bi, S. G. Schroeder, M. Yamasaki, J. F. Doebley, et al., 2005 The effects of artificial selection on the maize genome. Science 308: 1310–1314. 10.1126/science.1107891

Yang J., W.-R. Li, F.-H. Lv, S.-G. He, S.-L. Tian, et al., 2016 Whole-Genome Sequencing of Native Sheep Provides Insights into Rapid Adaptations to Extreme Environments. Mol. Biol. Evol. 33: 2576–2592. 10.1093/molbev/msw129

Zeder M. A., 2012 The Domestication of Animals. J. Anthropol. Res. 68: 161–190. 10.3998/jar.0521004.0068.201

Zeder M. A., 2015 Core questions in domestication research. Proc. Natl. Acad. Sci. 112: 3191–3198. 10.1073/pnas.1501711112

Zeder M. A., E. Emshwiller, B. D. Smith, and D. G. Bradley, 2006 Documenting domestication: the intersection of genetics and archaeology. Trends Genet. TIG 22: 139–155. 10.1016/j.tig.2006.01.007

Zeng, K., and B. Charlesworth, 2011 The joint effects of background selection and genetic recombination on local gene genealogies. Genetics 189: 251–266.

