## Supplemental Figures and Tables for "Detection of Domestication Signals through the Analysis of the Full Distribution of Fitness Effects"

### Diagnostic plots 2D-SFS

Synonymous data

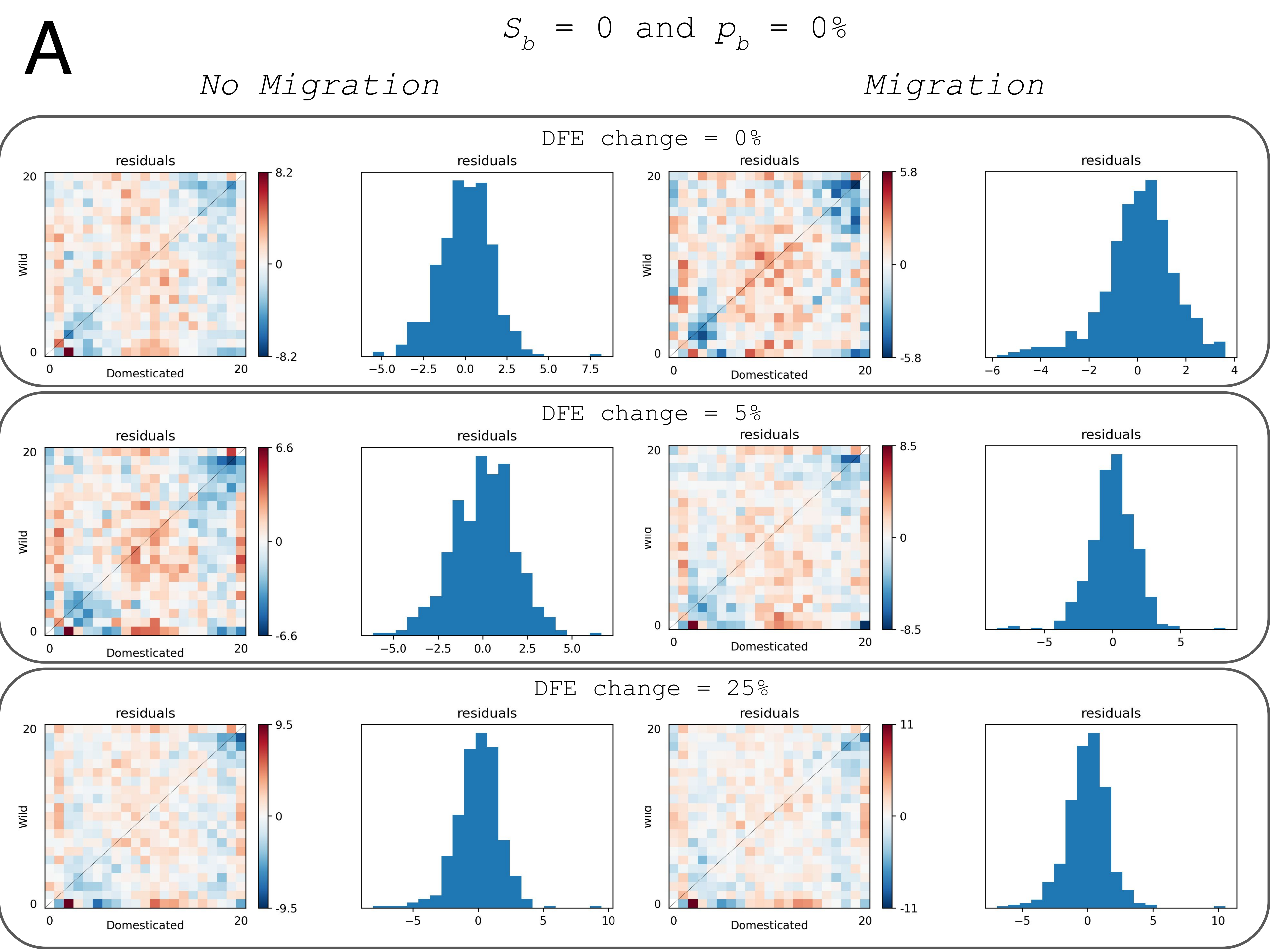

A
$S_b$ = 0 and $p_b$ = 0%
No Migration
Migration
DFE change = 0%
DFE change = 5%
DFE change = 25%
residuals
Wild
Domesticated

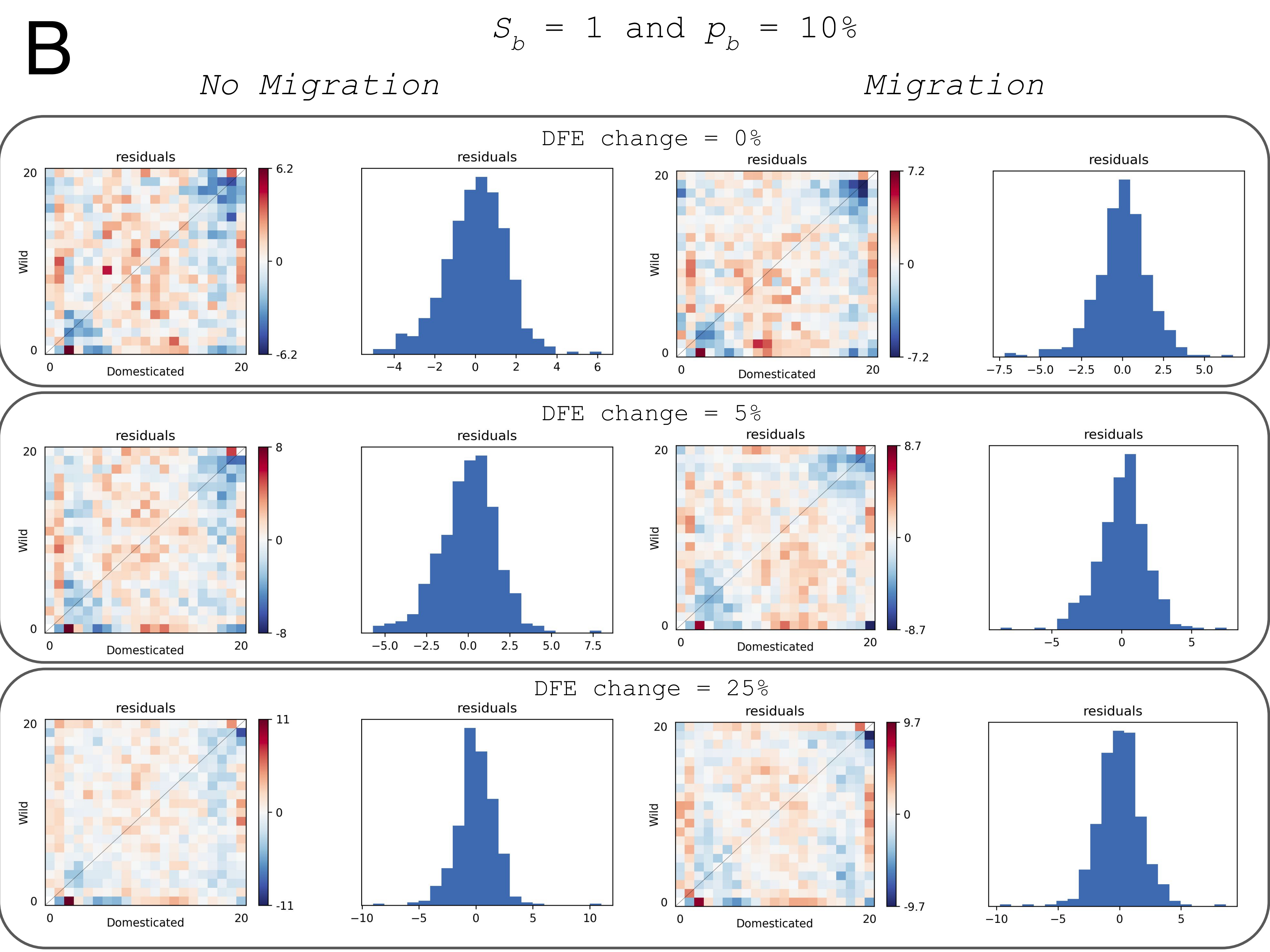
B
$S_b$ = 1 and $p_b$ = 10%
No Migration
Migration
DFE change = 0%
DFE change = 5%
DFE change = 25%
residuals
Wild
Domesticated

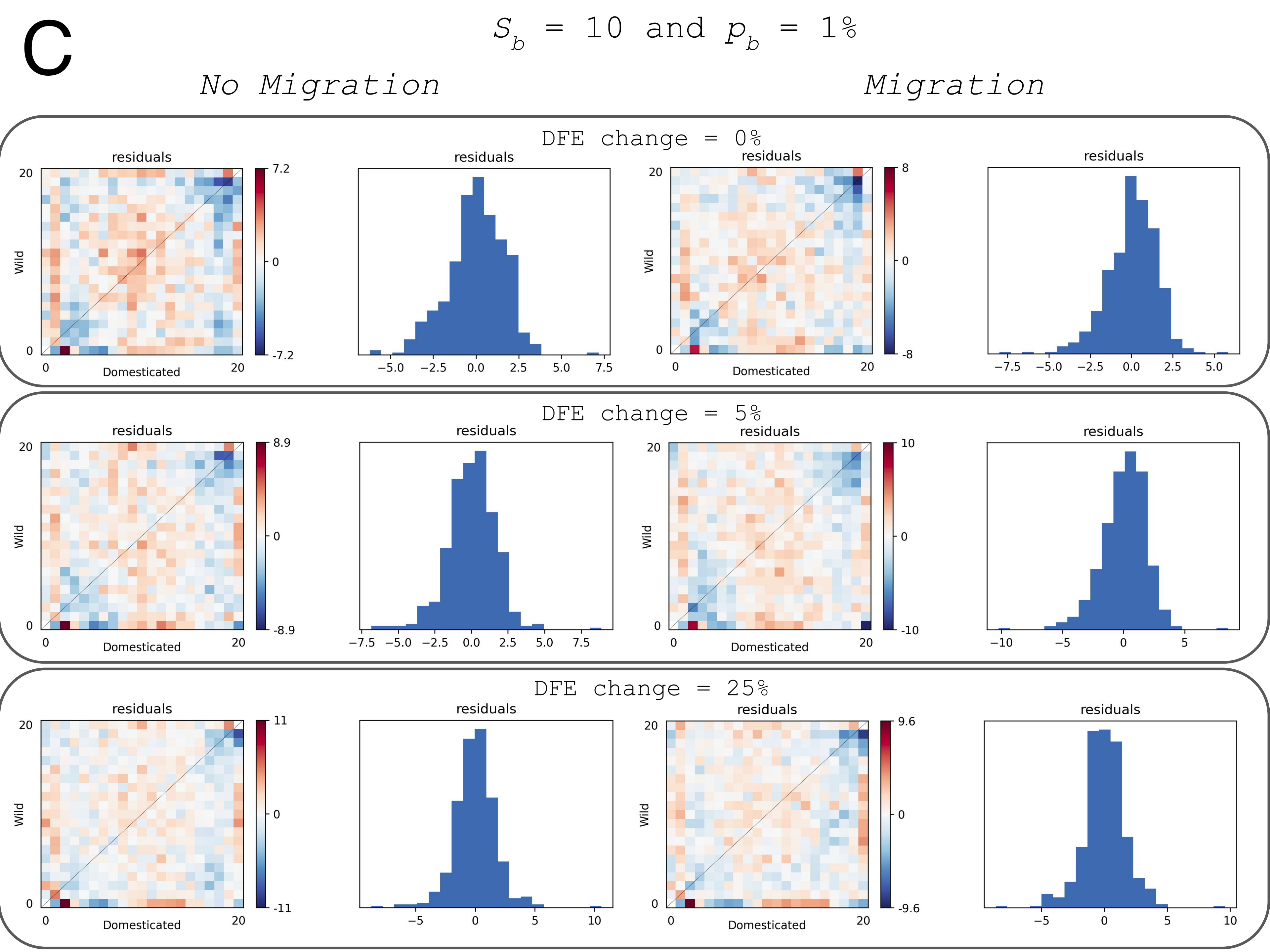

C
$S_b$ = 10 and $p_b$ = 1%
No Migration
Migration
DFE change = 0%
DFE change = 5%
DFE change = 25%
residuals
Wild
Domesticated

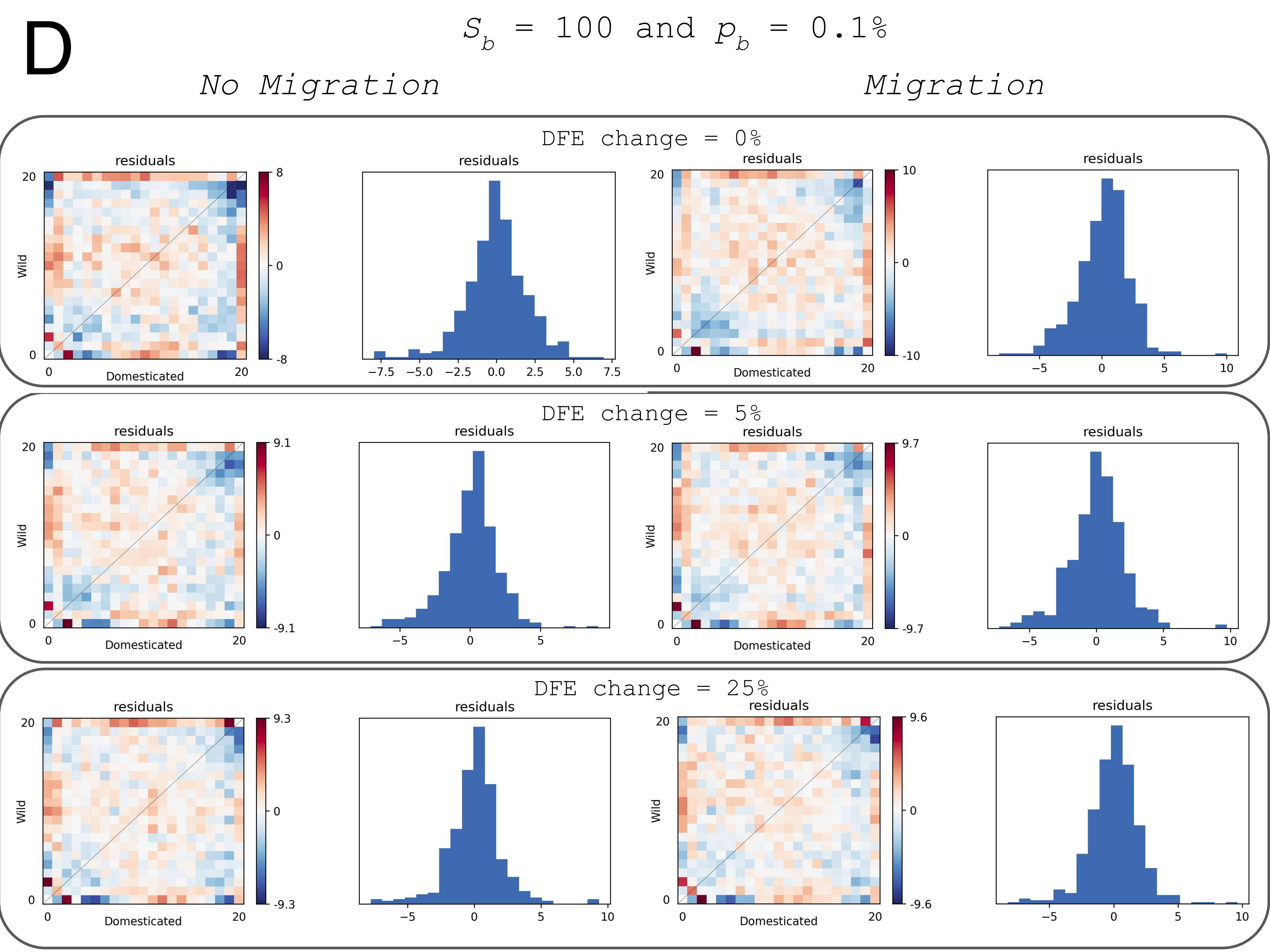

D
$S_b$ = 100 and $p_b$ = 0.1%
No Migration
Migration
DFE change = 0%
DFE change = 5%
DFE change = 25%
residuals
Wild
Domesticated
0
20
8
-8
10
-10
9.1
-9.1
9.7
-9.7
9.3
-9.3
9.6
-9.6
−7.5
−5.0
−2.5
0.0
2.5
5.0
7.5
−5
5
10

### Diagnostic plots 2D-SFS

Non-synonymous data

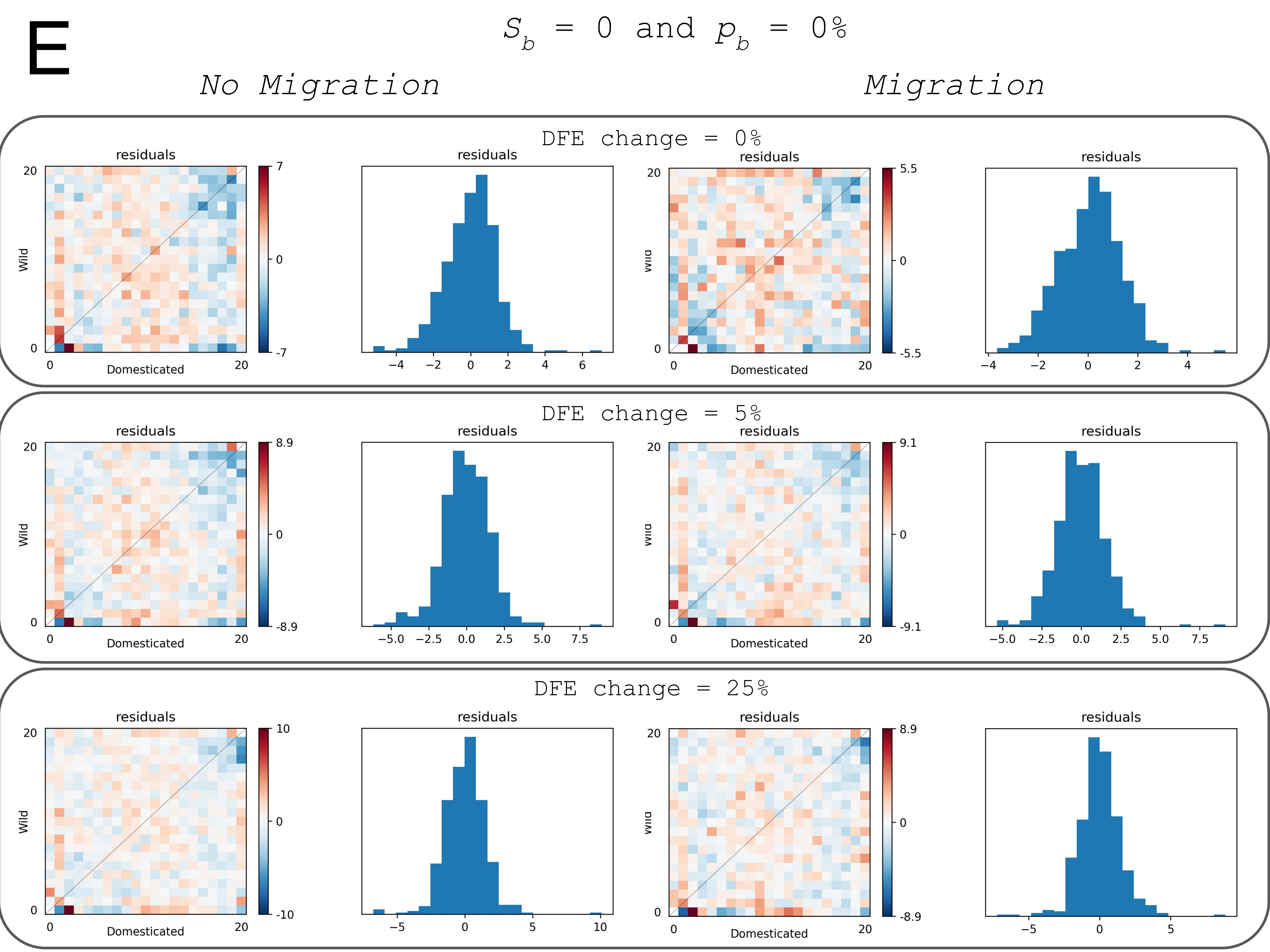
E
$S_b$ = 0 and $p_b$ = 0%
No Migration
Migration
DFE change = 0%
DFE change = 5%
DFE change = 25%
residuals
Wild
Domesticated

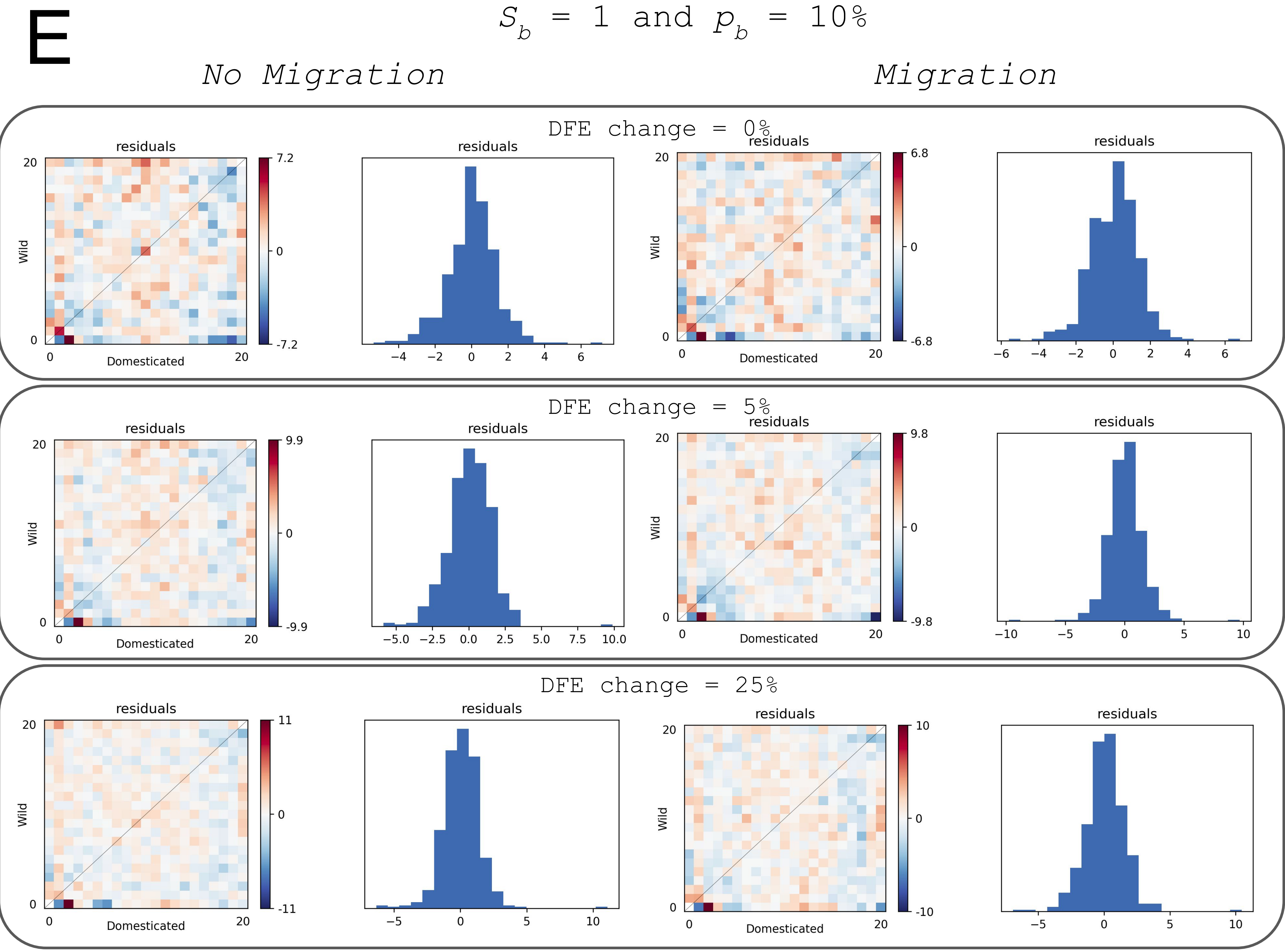

E
S_b = 1 and p_b = 10%
No Migration
Migration
DFE change = 0%
DFE change = 5%
DFE change = 25%
residuals
Wild
Domesticated

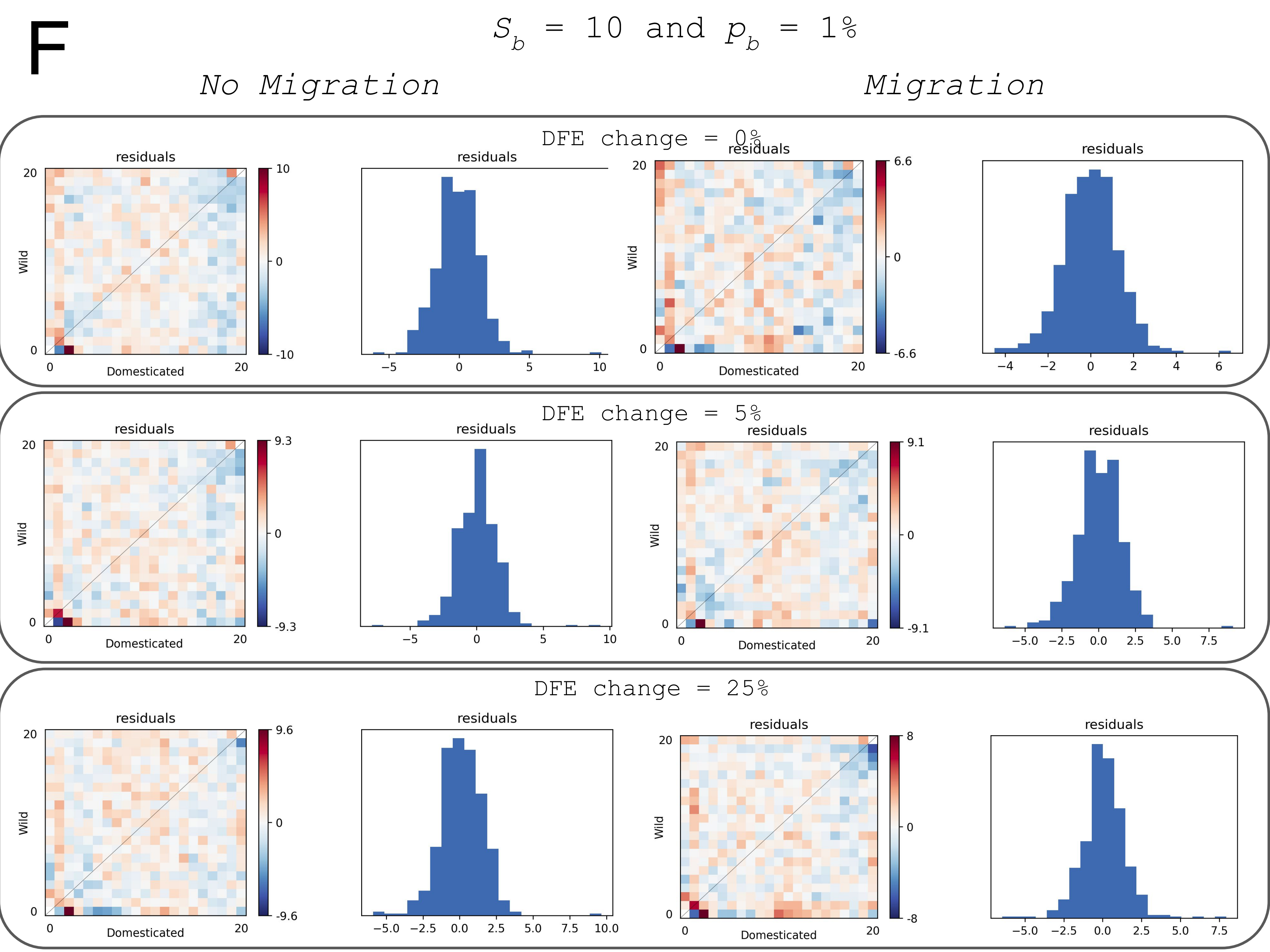

F
$S_b$ = 10 and $p_b$ = 1%
No Migration
Migration
DFE change = 0%
DFE change = 5%
DFE change = 25%
residuals
Wild
Domesticated

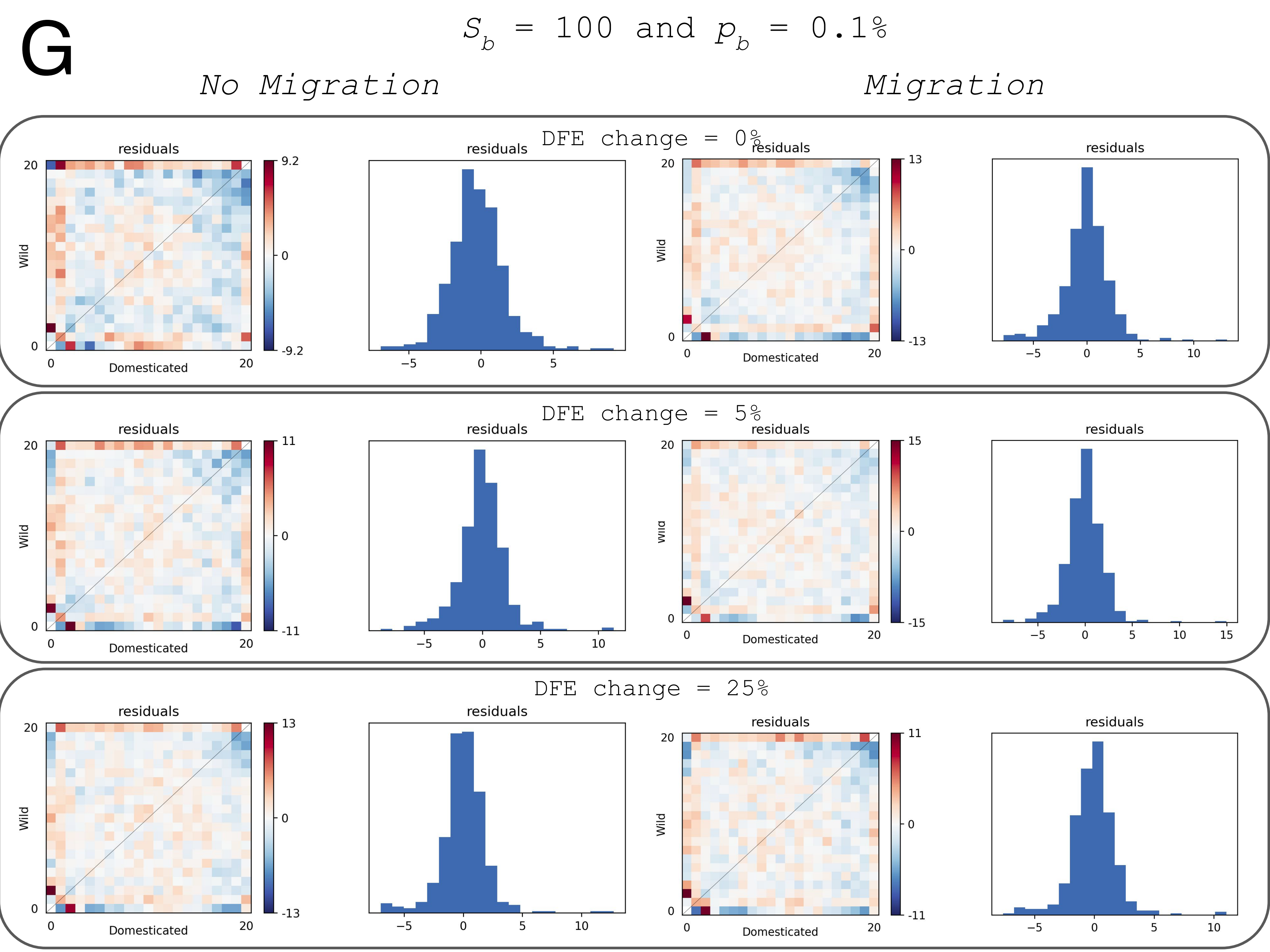

G
$S_b$ = 100 and $p_b$ = 0.1%
No Migration
Migration
DFE change = 0%
DFE change = 5%
DFE change = 25%
residuals
Wild
Domesticated

**Supplementary Figure 1.** The first four subfigures (A,B,C,D) show the concordance of the simulated data versus the inferred data for synonymous positions. (A) Results for the scenario with the parameters Sb=0, pb=0%, the subfigure A is separated in three rows (first for the scenarios having no change of mutation effects, then 5% change and last 25%). Each row shows the residuals (indicating in colors the difference between the 2D-SFS simulated scenario and the inference performed with dadi) plus the histogram of these residuals, in scenarios without (left) and with (right) migration (see Table 1 for all scenarios). (B) Same than A but for the scenario with the parameters Sb=1, pb=10%. (C) Same than A but for the scenario with the parameters Sb=100, pb=1%. (D) Same than A but for the scenario with the parameters Sb=100, pb=0.1%. Subfigures E, F, G and H show the same than A to D but considering nonsynonymous data.

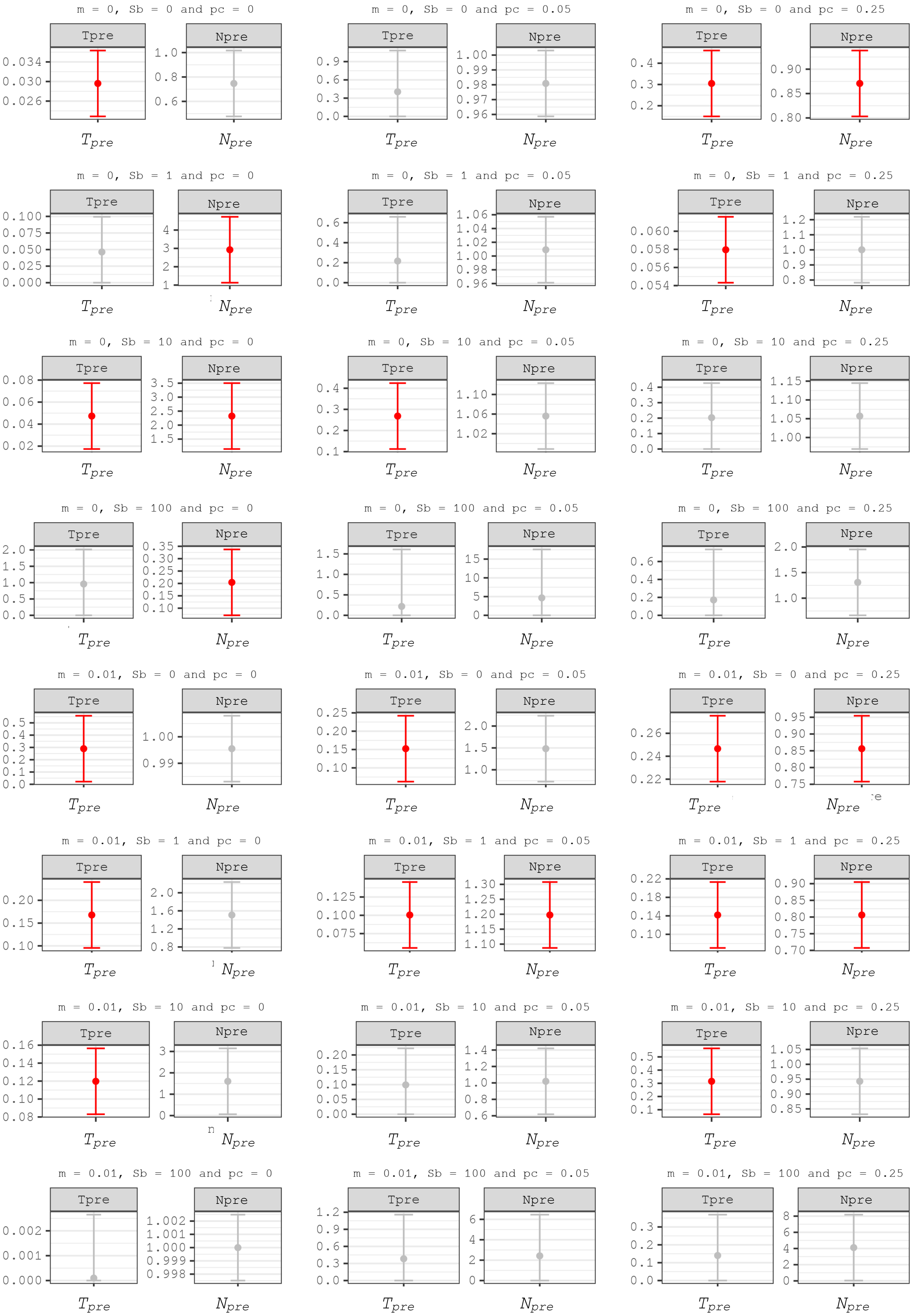
m = 0, Sb = 0 and pc = 0
m = 0, Sb = 0 and pc = 0.05
m = 0, Sb = 0 and pc = 0.25
m = 0, Sb = 1 and pc = 0
m = 0, Sb = 1 and pc = 0.05
m = 0, Sb = 1 and pc = 0.25
m = 0, Sb = 10 and pc = 0
m = 0, Sb = 10 and pc = 0.05
m = 0, Sb = 10 and pc = 0.25
m = 0, Sb = 100 and pc = 0
m = 0, Sb = 100 and pc = 0.05
m = 0, Sb = 100 and pc = 0.25
m = 0.01, Sb = 0 and pc = 0
m = 0.01, Sb = 0 and pc = 0.05
m = 0.01, Sb = 0 and pc = 0.25
m = 0.01, Sb = 1 and pc = 0
m = 0.01, Sb = 1 and pc = 0.05
m = 0.01, Sb = 1 and pc = 0.25
m = 0.01, Sb = 10 and pc = 0
m = 0.01, Sb = 10 and pc = 0.05
m = 0.01, Sb = 10 and pc = 0.25
m = 0.01, Sb = 100 and pc = 0
m = 0.01, Sb = 100 and pc = 0.05
m = 0.01, Sb = 100 and pc = 0.25
Tpre
Npre
$T_{pre}$
$N_{pre}$

**Supplementary Figure 2**. Confidence intervals of the population size change ($N_{pre}$) and time ($T_{pre}$) before the domestication split for all simulated scenarios. In red significant intervals, in grey intervals that overlap 0 ($T_{pre}$) or 1 ($N_{pre}$). A population size change *prior* to domestication must be considered valid only when both parameters are highlighted in red.

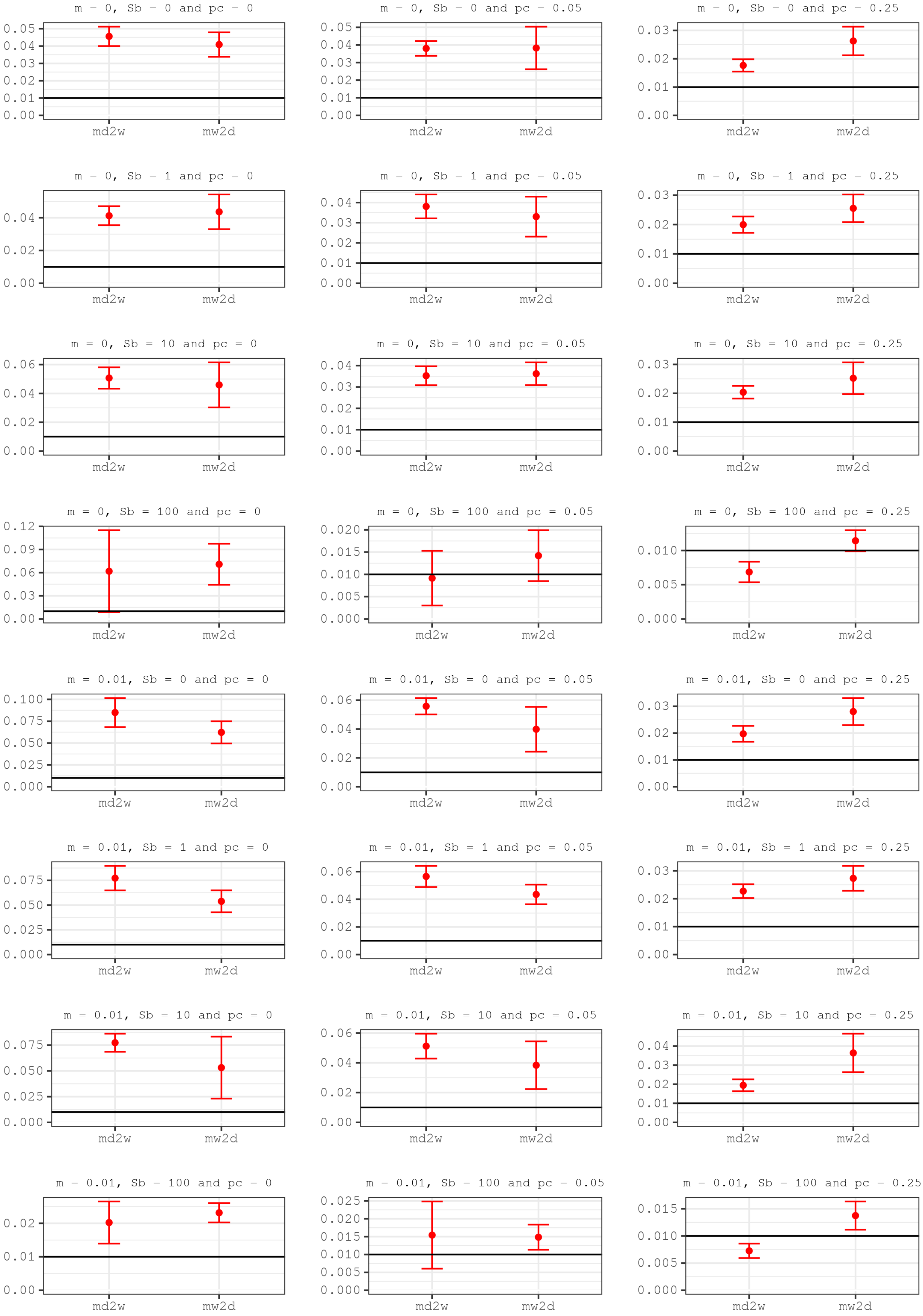
m = 0, Sb = 0 and pc = 0
m = 0, Sb = 0 and pc = 0.05
m = 0, Sb = 0 and pc = 0.25
m = 0, Sb = 1 and pc = 0
m = 0, Sb = 1 and pc = 0.05
m = 0, Sb = 1 and pc = 0.25
m = 0, Sb = 10 and pc = 0
m = 0, Sb = 10 and pc = 0.05
m = 0, Sb = 10 and pc = 0.25
m = 0, Sb = 100 and pc = 0
m = 0, Sb = 100 and pc = 0.05
m = 0, Sb = 100 and pc = 0.25
m = 0.01, Sb = 0 and pc = 0
m = 0.01, Sb = 0 and pc = 0.05
m = 0.01, Sb = 0 and pc = 0.25
m = 0.01, Sb = 1 and pc = 0
m = 0.01, Sb = 1 and pc = 0.05
m = 0.01, Sb = 1 and pc = 0.25
m = 0.01, Sb = 10 and pc = 0
m = 0.01, Sb = 10 and pc = 0.05
m = 0.01, Sb = 10 and pc = 0.25
m = 0.01, Sb = 100 and pc = 0
m = 0.01, Sb = 100 and pc = 0.05
m = 0.01, Sb = 100 and pc = 0.25
md2w
mw2d

**Supplementary Figure 3**. Confidence intervals the migration rate from Domestic to Wild ($m_{d2w}$) and from Wild to Domestic ($m_{w2d}$) for all simulated scenarios. In red significant intervals, in grey intervals that overlap 0. The horizontal solid line shows the simulated $m_{w2d}$. The simulated $m_{d2w}$ is 0.

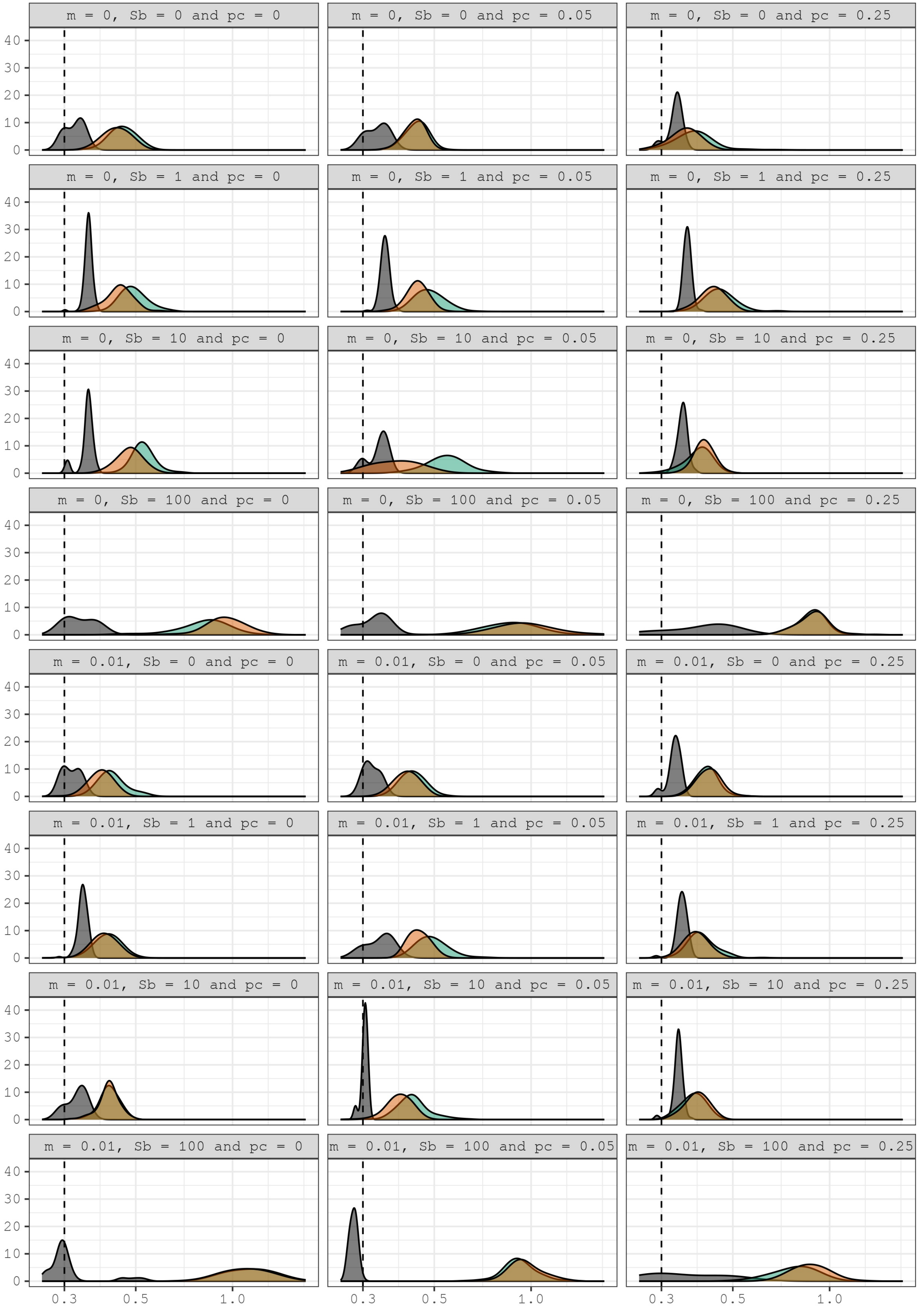
m = 0, Sb = 0 and pc = 0
m = 0, Sb = 0 and pc = 0.05
m = 0, Sb = 0 and pc = 0.25
m = 0, Sb = 1 and pc = 0
m = 0, Sb = 1 and pc = 0.05
m = 0, Sb = 1 and pc = 0.25
m = 0, Sb = 10 and pc = 0
m = 0, Sb = 10 and pc = 0.05
m = 0, Sb = 10 and pc = 0.25
m = 0, Sb = 100 and pc = 0
m = 0, Sb = 100 and pc = 0.05
m = 0, Sb = 100 and pc = 0.25
m = 0.01, Sb = 0 and pc = 0
m = 0.01, Sb = 0 and pc = 0.05
m = 0.01, Sb = 0 and pc = 0.25
m = 0.01, Sb = 1 and pc = 0
m = 0.01, Sb = 1 and pc = 0.05
m = 0.01, Sb = 1 and pc = 0.25
m = 0.01, Sb = 10 and pc = 0
m = 0.01, Sb = 10 and pc = 0.05
m = 0.01, Sb = 10 and pc = 0.25
m = 0.01, Sb = 100 and pc = 0
m = 0.01, Sb = 100 and pc = 0.05
m = 0.01, Sb = 100 and pc = 0.25
0
10
20
30
40
0.3
0.5
1.0

**Supplementary Figure 4**. Shape parameter distribution for each of the 24 different model conditions estimated with dadi. (grey) and polyDFE (turquoise green and salmon-orange). The conditions are indicated above each representation. The ancestral population is represented in grey, in turquoise green the domestic population and in salmon-orange the wild population. Sampling distributions of estimated parameters for the deleterious DFE are obtained using 100 bootstrap replicates. Dotted vertical lines indicate the actual simulated parameter value.

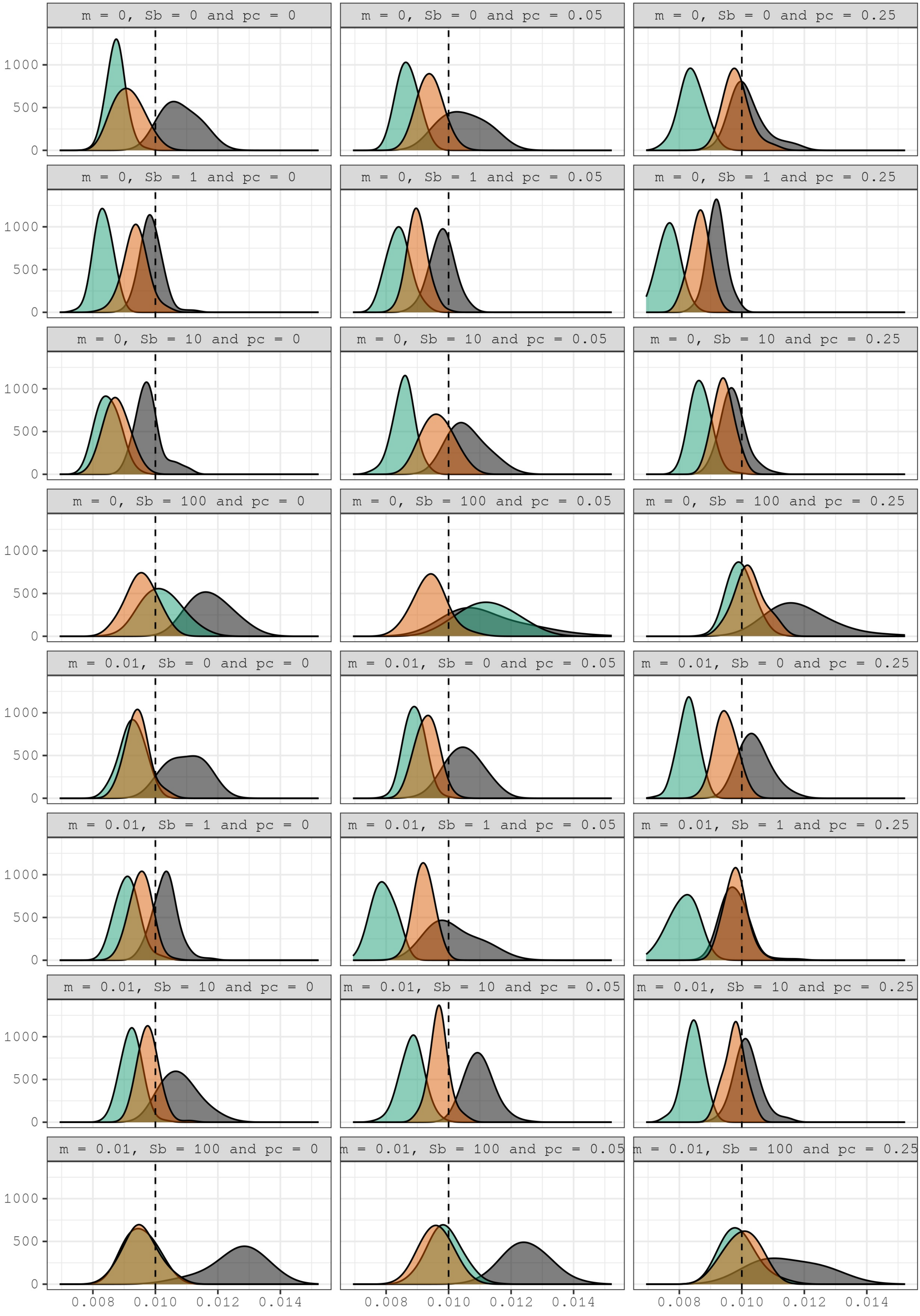

m = 0, Sb = 0 and pc = 0
m = 0, Sb = 0 and pc = 0.05
m = 0, Sb = 0 and pc = 0.25
m = 0, Sb = 1 and pc = 0
m = 0, Sb = 1 and pc = 0.05
m = 0, Sb = 1 and pc = 0.25
m = 0, Sb = 10 and pc = 0
m = 0, Sb = 10 and pc = 0.05
m = 0, Sb = 10 and pc = 0.25
m = 0, Sb = 100 and pc = 0
m = 0, Sb = 100 and pc = 0.05
m = 0, Sb = 100 and pc = 0.25
m = 0.01, Sb = 0 and pc = 0
m = 0.01, Sb = 0 and pc = 0.05
m = 0.01, Sb = 0 and pc = 0.25
m = 0.01, Sb = 1 and pc = 0
m = 0.01, Sb = 1 and pc = 0.05
m = 0.01, Sb = 1 and pc = 0.25
m = 0.01, Sb = 10 and pc = 0
m = 0.01, Sb = 10 and pc = 0.05
m = 0.01, Sb = 10 and pc = 0.25
m = 0.01, Sb = 100 and pc = 0
m = 0.01, Sb = 100 and pc = 0.05
m = 0.01, Sb = 100 and pc = 0.25
0
500
1000
0.008
0.010
0.012
0.014

**Supplementary Figure 5.** mean *S* estimated with dadi. (grey) and polyDFE (turquoise green and salmon-orange). for each of the twenty-four different model conditions. The ancestral population is represented in grey, in turquoise green the domestic population and in salmon-orange the wild population. To calculate *s* from inferred *S* values, we divided *S* by 4 times the $N_e$ estimate in polyDFE and by 2 times the $N_a$ estimate in dadi. To obtain the $N_a$ we divide π at synonymous sites by the true simulated mutation rate (2.5x10-7 per site and generation).

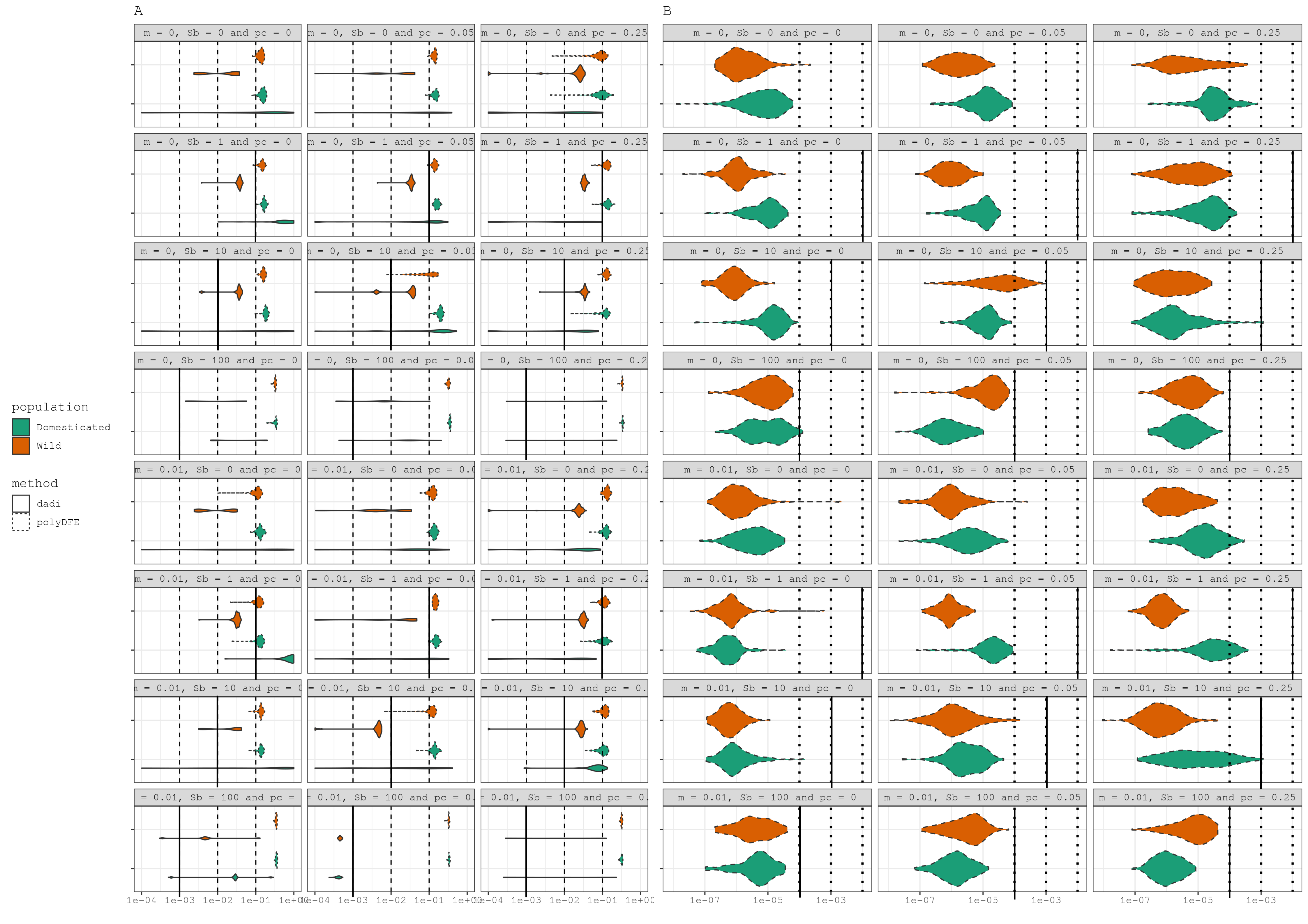
A
B
population
Domesticated
Wild
method
dadi
polyDFE
m = 0, Sb = 0 and pc = 0
m = 0, Sb = 0 and pc = 0.05
m = 0, Sb = 0 and pc = 0.25
m = 0, Sb = 1 and pc = 0
m = 0, Sb = 1 and pc = 0.05
m = 0, Sb = 1 and pc = 0.25
m = 0, Sb = 10 and pc = 0
m = 0, Sb = 10 and pc = 0.05
m = 0, Sb = 10 and pc = 0.25
m = 0, Sb = 100 and pc = 0
m = 0, Sb = 100 and pc = 0.05
m = 0, Sb = 100 and pc = 0.25
m = 0.01, Sb = 0 and pc = 0
m = 0.01, Sb = 0 and pc = 0.05
m = 0.01, Sb = 0 and pc = 0.25
m = 0.01, Sb = 1 and pc = 0
m = 0.01, Sb = 1 and pc = 0.05
m = 0.01, Sb = 1 and pc = 0.25
m = 0.01, Sb = 10 and pc = 0
m = 0.01, Sb = 10 and pc = 0.05
m = 0.01, Sb = 10 and pc = 0.25
m = 0.01, Sb = 100 and pc = 0
m = 0.01, Sb = 100 and pc = 0.05
m = 0.01, Sb = 100 and pc = 0.25
1e-04 1e-03 1e-02 1e-01 1e+00
1e-07 1e-05 1e-03

**Supplementary Figure 6.** Sampling distributions of estimated parameters for the beneficial DFE are obtained using 100 bootstrap replicates. Dotted vertical lines indicate the actual simulated parameter values. Orange color indicates the Wild population while green color indicates the Domestic population. Violin distribution plots with a dotted contour refer to polyDFE method while a continuous contour refer to dadi method . A) $p_b$ parameter estimations for wild and Domestic populations inferred with polyDFE and dadi methods for the twenty-four simulated scenarios, B) mean $s_b$ estimations for wild and Domestic populations inferred with polyDFE methods for the twenty-four simulated scenarios (note that with *dadi* we must assume a given point mass distribution for the beneficial DFE for computational reasons). To obtain $N_e$ we divide π at synonymous sites by the true simulated mutation rate (2.5x10-7 per site and generation). Supplementary table 3 shows the most likely $S_b$ values with *dadi*.

A

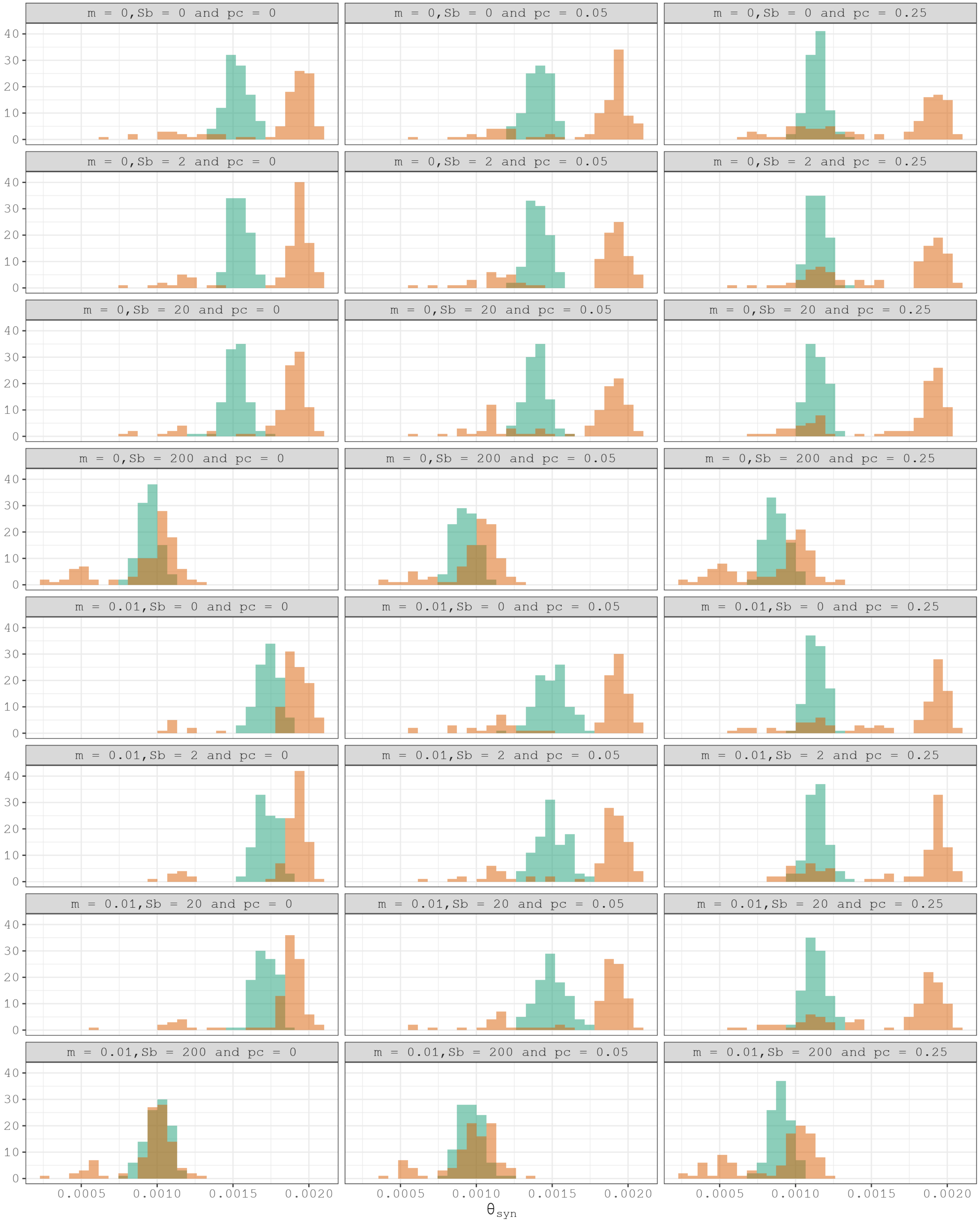
m = 0,Sb = 0 and pc = 0
m = 0,Sb = 0 and pc = 0.05
m = 0,Sb = 0 and pc = 0.25
m = 0,Sb = 2 and pc = 0
m = 0,Sb = 2 and pc = 0.05
m = 0,Sb = 2 and pc = 0.25
m = 0,Sb = 20 and pc = 0
m = 0,Sb = 20 and pc = 0.05
m = 0,Sb = 20 and pc = 0.25
m = 0,Sb = 200 and pc = 0
m = 0,Sb = 200 and pc = 0.05
m = 0,Sb = 200 and pc = 0.25
m = 0.01,Sb = 0 and pc = 0
m = 0.01,Sb = 0 and pc = 0.05
m = 0.01,Sb = 0 and pc = 0.25
m = 0.01,Sb = 2 and pc = 0
m = 0.01,Sb = 2 and pc = 0.05
m = 0.01,Sb = 2 and pc = 0.25
m = 0.01,Sb = 20 and pc = 0
m = 0.01,Sb = 20 and pc = 0.05
m = 0.01,Sb = 20 and pc = 0.25
m = 0.01,Sb = 200 and pc = 0
m = 0.01,Sb = 200 and pc = 0.05
m = 0.01,Sb = 200 and pc = 0.25
0
10
20
30
40
0.0005
0.0010
0.0015
0.0020
$\theta_{syn}$

B

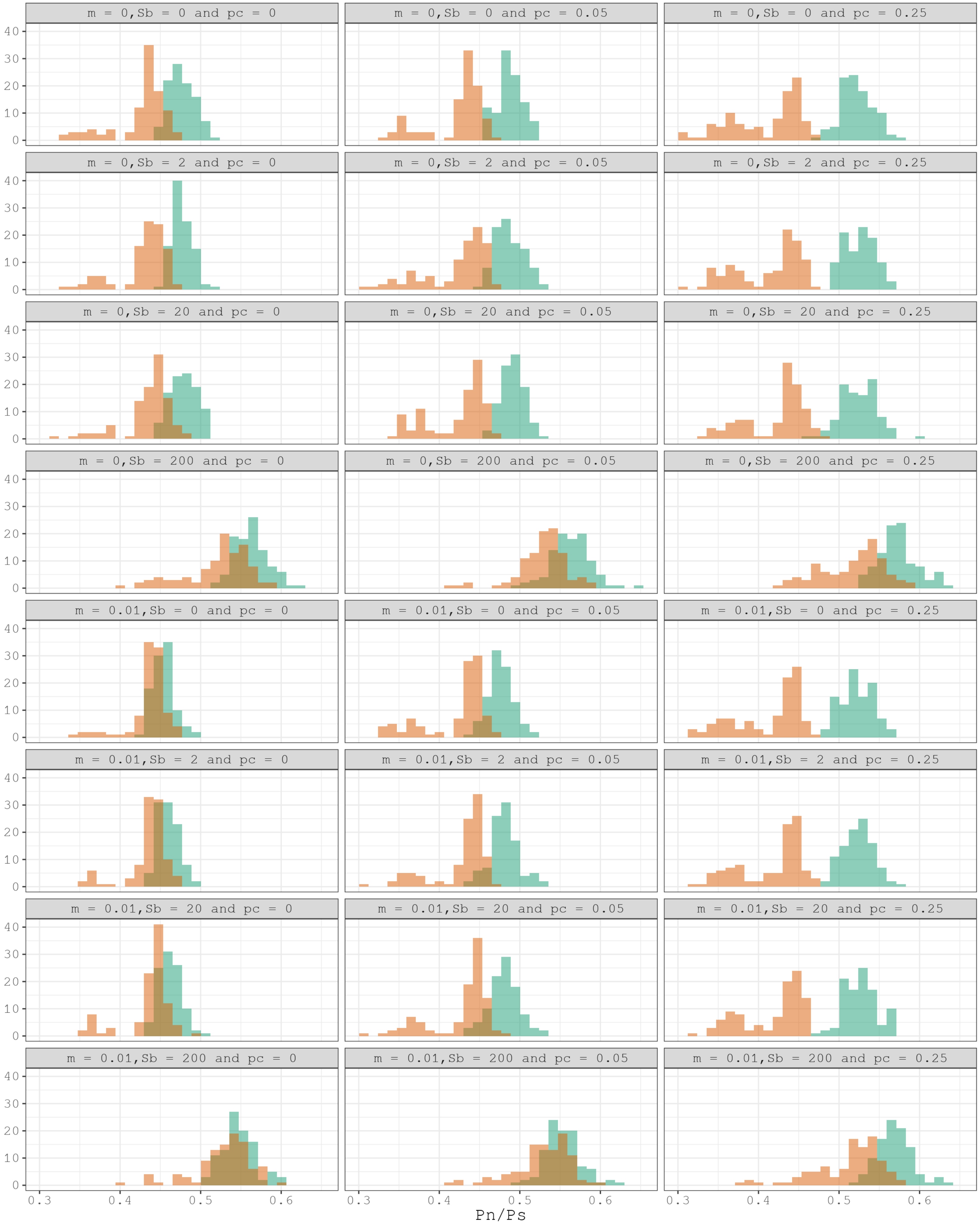
m = 0, Sb = 0 and pc = 0
m = 0, Sb = 0 and pc = 0.05
m = 0, Sb = 0 and pc = 0.25
m = 0, Sb = 2 and pc = 0
m = 0, Sb = 2 and pc = 0.05
m = 0, Sb = 2 and pc = 0.25
m = 0, Sb = 20 and pc = 0
m = 0, Sb = 20 and pc = 0.05
m = 0, Sb = 20 and pc = 0.25
m = 0, Sb = 200 and pc = 0
m = 0, Sb = 200 and pc = 0.05
m = 0, Sb = 200 and pc = 0.25
m = 0.01, Sb = 0 and pc = 0
m = 0.01, Sb = 0 and pc = 0.05
m = 0.01, Sb = 0 and pc = 0.25
m = 0.01, Sb = 2 and pc = 0
m = 0.01, Sb = 2 and pc = 0.05
m = 0.01, Sb = 2 and pc = 0.25
m = 0.01, Sb = 20 and pc = 0
m = 0.01, Sb = 20 and pc = 0.05
m = 0.01, Sb = 20 and pc = 0.25
m = 0.01, Sb = 200 and pc = 0
m = 0.01, Sb = 200 and pc = 0.05
m = 0.01, Sb = 200 and pc = 0.25
0
10
20
30
40
0.3
0.4
0.5
0.6
Pn/Ps

**Supplementary Figure 7.** A) Histogram of neutral genetic diversity (*θs*). B) Histogram of selective constraint (*Pn/Ps*) across the twenty-four domestication scenarios. Each data point is an independent simulation run. In salmon-orange the Wild population and in turquoise-green the Domestic population.

| | Scenario 1 | | | Scenario 2 | | | Scenario 3 | | | Scenario 4 | | | Scenario 5 | | | Scenario 6 | | |
|---|---|---|---|---|---|---|---|---|---|---|---|---|---|---|---|---|---|---|
| **Parameters*** | **CI_2.5%** | **CI_97.5%** | **value** | **CI_2.5%** | **CI_97.5%** | **value** | **CI_2.5%** | **CI_97.5%** | **value** | **CI_2.5%** | **CI_97.5%** | **value** | **CI_2.5%** | **CI_97.5%** | **value** | **CI_2.5%** | **CI_97.5%** | **value** |
| ***Tpre*** | 0.023 | 0.036 | 0.030 | 0.000 | 1.083 | 0.404 | 0.150 | 0.460 | 0.305 | 0.000 | 0.099 | 0.046 | 0.000 | 0.659 | 0.217 | 0.054 | 0.062 | 0.058 |
| ***nuPre*** | 0.477 | 1.017 | 0.747 | 0.959 | 1.003 | 0.981 | 0.803 | 0.938 | 0.871 | 1.129 | 4.721 | 2.925 | 0.961 | 1.057 | 1.009 | 0.782 | 1.219 | 1.001 |
| ***Tdiv*** | 0.133 | 0.139 | 0.136 | 0.118 | 0.142 | 0.130 | 0.140 | 0.172 | 0.156 | 0.039 | 0.162 | 0.101 | 0.103 | 0.148 | 0.126 | 0.126 | 0.135 | 0.130 |
| ***nu1div*** | 1.148 | 1.255 | 1.202 | 0.922 | 1.189 | 1.055 | 0.876 | 1.056 | 0.966 | 0.744 | 1.140 | 0.942 | 0.687 | 1.304 | 0.996 | 0.897 | 0.998 | 0.947 |
| ***nu2div*** | 0.207 | 0.214 | 0.210 | 0.132 | 0.163 | 0.147 | 0.084 | 0.103 | 0.093 | 0.074 | 0.266 | 0.170 | 0.119 | 0.169 | 0.144 | 0.074 | 0.079 | 0.076 |
| ***T1F*** | 0.102 | 0.111 | 0.107 | 0.109 | 0.206 | 0.157 | 0.168 | 0.200 | 0.184 | 0.126 | 0.210 | 0.168 | 0.050 | 0.285 | 0.167 | 0.157 | 0.182 | 0.170 |
| ***T2F*** | 0.240 | 0.257 | 0.248 | 0.255 | 0.273 | 0.264 | 0.231 | 0.249 | 0.240 | 0.229 | 0.295 | 0.262 | 0.247 | 0.266 | 0.257 | 0.249 | 0.259 | 0.254 |
| ***nu1F*** | 2.200 | 2.377 | 2.288 | 1.933 | 2.363 | 2.148 | 1.914 | 2.039 | 1.976 | 2.023 | 2.314 | 2.168 | 1.732 | 2.481 | 2.106 | 2.002 | 2.182 | 2.092 |
| ***nu2F*** | 1.836 | 1.970 | 1.903 | 1.811 | 1.913 | 1.862 | 1.659 | 1.849 | 1.754 | 1.809 | 1.980 | 1.895 | 1.774 | 1.878 | 1.826 | 1.712 | 1.791 | 1.752 |
| ***mw2d*** | 0.034 | 0.048 | 0.041 | 0.026 | 0.050 | 0.038 | 0.021 | 0.031 | 0.026 | 0.033 | 0.054 | 0.044 | 0.023 | 0.043 | 0.033 | 0.021 | 0.030 | 0.026 |
| ***md2w*** | 0.040 | 0.051 | 0.046 | 0.034 | 0.042 | 0.038 | 0.015 | 0.020 | 0.018 | 0.035 | 0.047 | 0.041 | 0.032 | 0.044 | 0.038 | 0.017 | 0.023 | 0.020 |
| ***missid*** | 0.003 | 0.005 | 0.004 | 0.003 | 0.005 | 0.004 | 0.001 | 0.002 | 0.001 | 0.003 | 0.005 | 0.004 | 0.003 | 0.004 | 0.004 | 0.001 | 0.002 | 0.002 |

| | Scenario 7 | | | Scenario 8 | | | Scenario 9 | | | Scenario 10 | | | Scenario 11 | | | Scenario 12 | | |
|---|---|---|---|---|---|---|---|---|---|---|---|---|---|---|---|---|---|---|
| **Parameters*** | **CI_2.5%** | **CI_97.5%** | **value** | **CI_2.5%** | **CI_97.5%** | **value** | **CI_2.5%** | **CI_97.5%** | **value** | **CI_2.5%** | **CI_97.5%** | **value** | **CI_2.5%** | **CI_97.5%** | **value** | **CI_2.5%** | **CI_97.5%** | **value** |
| ***Tpre*** | 0.017 | 0.077 | 0.047 | 0.112 | 0.424 | 0.268 | 0.000 | 0.426 | 0.203 | 0.000 | 2.016 | 0.958 | 0.000 | 1.609 | 0.218 | 0.000 | 0.735 | 0.171 |
| ***nuPre*** | 1.145 | 3.502 | 2.323 | 0.989 | 1.123 | 1.056 | 0.969 | 1.145 | 1.057 | 0.071 | 0.337 | 0.204 | 0.000 | 17.580 | 4.656 | 0.665 | 1.952 | 1.309 |
| ***Tdiv*** | 0.078 | 0.150 | 0.114 | 0.128 | 0.134 | 0.131 | 0.137 | 0.155 | 0.146 | 0.068 | 0.187 | 0.127 | 0.000 | 1.332 | 0.472 | 0.464 | 0.705 | 0.584 |
| ***nu1div*** | 0.774 | 1.140 | 0.957 | 1.009 | 1.093 | 1.051 | 0.801 | 0.889 | 0.845 | 0.088 | 0.254 | 0.171 | 0.000 | 2.045 | 0.686 | 0.595 | 0.850 | 0.722 |
| ***nu2div*** | 0.124 | 0.228 | 0.176 | 0.145 | 0.151 | 0.148 | 0.080 | 0.092 | 0.086 | 0.062 | 0.177 | 0.120 | 0.000 | 1.003 | 0.400 | 0.333 | 0.399 | 0.366 |
| ***T1F*** | 0.115 | 0.236 | 0.175 | 0.150 | 0.153 | 0.151 | 0.225 | 0.226 | 0.225 | 0.041 | 0.142 | 0.091 | 0.000 | 1.211 | 0.503 | 0.350 | 0.579 | 0.464 |
| ***T2F*** | 0.250 | 0.285 | 0.268 | 0.267 | 0.281 | 0.274 | 0.250 | 0.261 | 0.255 | 0.036 | 0.139 | 0.088 | 0.061 | 0.977 | 0.519 | 0.393 | 0.524 | 0.458 |
| ***nu1F*** | 1.975 | 2.423 | 2.199 | 2.249 | 2.485 | 2.367 | 1.990 | 2.074 | 2.032 | 0.495 | 1.507 | 1.001 | 4.047 | 5.807 | 4.927 | 4.387 | 4.823 | 4.605 |
| ***nu2F*** | 1.900 | 2.020 | 1.960 | 1.872 | 1.913 | 1.893 | 1.816 | 1.820 | 1.818 | 0.532 | 1.517 | 1.025 | 4.380 | 4.905 | 4.642 | 4.094 | 4.732 | 4.413 |
| ***mw2d*** | 0.030 | 0.062 | 0.046 | 0.031 | 0.042 | 0.036 | 0.020 | 0.031 | 0.025 | 0.044 | 0.097 | 0.071 | 0.008 | 0.020 | 0.014 | 0.010 | 0.013 | 0.011 |
| ***md2w*** | 0.043 | 0.058 | 0.051 | 0.031 | 0.040 | 0.035 | 0.018 | 0.023 | 0.020 | 0.009 | 0.115 | 0.062 | 0.003 | 0.015 | 0.009 | 0.005 | 0.008 | 0.007 |
| ***missid*** | 0.003 | 0.005 | 0.004 | 0.003 | 0.004 | 0.004 | 0.001 | 0.002 | 0.002 | 0.001 | 0.005 | 0.003 | 0.002 | 0.006 | 0.004 | 0.002 | 0.005 | 0.004 |

| | Scenario 13 | | | Scenario 14 | | | Scenario 15 | | | Scenario 16 | | | Scenario 17 | | | Scenario 18 | | |
|---|---|---|---|---|---|---|---|---|---|---|---|---|---|---|---|---|---|---|
| **Parameters*** | **CI_2.5%** | **CI_97.5%** | **value** | **CI_2.5%** | **CI_97.5%** | **value** | **CI_2.5%** | **CI_97.5%** | **value** | **CI_2.5%** | **CI_97.5%** | **value** | **CI_2.5%** | **CI_97.5%** | **value** | **CI_2.5%** | **CI_97.5%** | **value** |
| ***Tpre*** | 0.022 | 0.558 | 0.290 | 0.062 | 0.242 | 0.152 | 0.218 | 0.275 | 0.247 | 0.095 | 0.240 | 0.168 | 0.055 | 0.145 | 0.100 | 0.071 | 0.213 | 0.142 |
| ***nuPre*** | 0.983 | 1.008 | 0.996 | 0.729 | 2.236 | 1.482 | 0.758 | 0.955 | 0.856 | 0.779 | 2.237 | 1.508 | 1.087 | 1.308 | 1.197 | 0.707 | 0.905 | 0.806 |
| ***Tdiv*** | 0.105 | 0.182 | 0.144 | 0.000 | 0.167 | 0.073 | 0.144 | 0.178 | 0.161 | 0.000 | 0.166 | 0.076 | 0.112 | 0.118 | 0.115 | 0.157 | 0.185 | 0.171 |
| ***nu1div*** | 1.081 | 1.249 | 1.165 | 0.457 | 1.053 | 0.755 | 1.018 | 1.194 | 1.106 | 0.701 | 1.043 | 0.872 | 0.894 | 1.010 | 0.952 | 1.050 | 1.137 | 1.093 |
| ***nu2div*** | 0.319 | 0.520 | 0.420 | 0.000 | 0.240 | 0.108 | 0.084 | 0.103 | 0.094 | 0.000 | 0.479 | 0.229 | 0.164 | 0.172 | 0.168 | 0.091 | 0.108 | 0.100 |
| ***T1F*** | 0.087 | 0.156 | 0.122 | 0.188 | 0.234 | 0.211 | 0.096 | 0.152 | 0.124 | 0.161 | 0.191 | 0.176 | 0.164 | 0.196 | 0.180 | 0.134 | 0.158 | 0.146 |
| ***T2F*** | 0.197 | 0.262 | 0.229 | 0.232 | 0.329 | 0.281 | 0.231 | 0.247 | 0.239 | 0.211 | 0.320 | 0.266 | 0.256 | 0.269 | 0.262 | 0.232 | 0.251 | 0.241 |
| ***nu1F*** | 2.010 | 2.426 | 2.218 | 1.958 | 2.255 | 2.107 | 2.090 | 2.329 | 2.210 | 2.116 | 2.345 | 2.230 | 2.047 | 2.203 | 2.125 | 2.056 | 2.234 | 2.145 |
| ***nu2F*** | 1.863 | 2.099 | 1.981 | 1.758 | 1.911 | 1.834 | 1.700 | 1.790 | 1.745 | 1.850 | 2.158 | 2.004 | 1.848 | 1.901 | 1.874 | 1.708 | 1.804 | 1.756 |
| ***mw2d*** | 0.050 | 0.075 | 0.062 | 0.024 | 0.055 | 0.040 | 0.023 | 0.033 | 0.028 | 0.043 | 0.065 | 0.054 | 0.036 | 0.051 | 0.043 | 0.023 | 0.032 | 0.027 |
| ***md2w*** | 0.068 | 0.102 | 0.085 | 0.050 | 0.062 | 0.056 | 0.017 | 0.023 | 0.020 | 0.065 | 0.090 | 0.077 | 0.049 | 0.064 | 0.057 | 0.020 | 0.025 | 0.023 |
| ***missid*** | 0.004 | 0.005 | 0.004 | 0.003 | 0.005 | 0.004 | 0.001 | 0.002 | 0.001 | 0.004 | 0.006 | 0.005 | 0.004 | 0.005 | 0.004 | 0.001 | 0.003 | 0.002 |

| | Scenario 19 | | | Scenario 20 | | | Scenario 21 | | | Scenario 22 | | | Scenario 23 | | | Scenario 24 | | |
|---|---|---|---|---|---|---|---|---|---|---|---|---|---|---|---|---|---|---|
| **Parameters*** | **CI_2.5%** | **CI_97.5%** | **value** | **CI_2.5%** | **CI_97.5%** | **value** | **CI_2.5%** | **CI_97.5%** | **value** | **CI_2.5%** | **CI_97.5%** | **value** | **CI_2.5%** | **CI_97.5%** | **value** | **CI_2.5%** | **CI_97.5%** | **value** |
| ***Tpre*** | 0.083 | 0.156 | 0.120 | 0.000 | 0.222 | 0.099 | 0.066 | 0.565 | 0.315 | 0.000 | 0.003 | 0.000 | 0.000 | 1.157 | 0.383 | 0.000 | 0.369 | 0.140 |
| ***nuPre*** | 0.062 | 3.145 | 1.604 | 0.619 | 1.420 | 1.020 | 0.832 | 1.054 | 0.943 | 0.998 | 1.002 | 1.000 | 0.000 | 6.460 | 2.422 | 0.040 | 8.182 | 4.111 |
| ***Tdiv*** | 0.000 | 0.224 | 0.097 | 0.071 | 0.162 | 0.117 | 0.144 | 0.183 | 0.164 | 0.000 | 1.449 | 0.442 | 0.000 | 1.002 | 0.370 | 0.409 | 0.626 | 0.517 |
| ***nu1div*** | 0.646 | 1.297 | 0.972 | 0.943 | 1.129 | 1.036 | 0.874 | 0.982 | 0.928 | 0.000 | 1.560 | 0.702 | 0.000 | 1.371 | 0.540 | 0.630 | 0.736 | 0.683 |
| ***nu2div*** | 0.000 | 0.643 | 0.295 | 0.109 | 0.235 | 0.172 | 0.085 | 0.110 | 0.097 | 0.000 | 1.750 | 0.552 | 0.158 | 0.748 | 0.453 | 0.289 | 0.403 | 0.346 |
| ***T1F*** | 0.042 | 0.306 | 0.174 | 0.156 | 0.162 | 0.159 | 0.194 | 0.194 | 0.194 | 0.386 | 0.453 | 0.420 | 0.032 | 1.053 | 0.543 | 0.449 | 0.498 | 0.473 |
| ***T2F*** | 0.189 | 0.343 | 0.266 | 0.245 | 0.294 | 0.269 | 0.233 | 0.257 | 0.245 | 0.132 | 0.802 | 0.467 | 0.257 | 0.720 | 0.489 | 0.420 | 0.548 | 0.484 |
| ***nu1F*** | 1.635 | 2.888 | 2.261 | 2.132 | 2.394 | 2.263 | 1.958 | 2.116 | 2.037 | 2.008 | 7.108 | 4.558 | 4.013 | 5.071 | 4.542 | 3.843 | 5.241 | 4.542 |
| ***nu2F*** | 1.947 | 2.157 | 2.052 | 1.849 | 1.934 | 1.891 | 1.791 | 1.795 | 1.793 | 0.000 | 8.560 | 4.242 | 4.001 | 5.412 | 4.706 | 3.987 | 4.773 | 4.380 |
| ***mw2d*** | 0.023 | 0.083 | 0.053 | 0.022 | 0.054 | 0.038 | 0.026 | 0.046 | 0.036 | 0.020 | 0.026 | 0.023 | 0.011 | 0.018 | 0.015 | 0.011 | 0.016 | 0.014 |
| ***md2w*** | 0.069 | 0.086 | 0.077 | 0.043 | 0.060 | 0.051 | 0.016 | 0.023 | 0.019 | 0.014 | 0.027 | 0.020 | 0.006 | 0.025 | 0.015 | 0.006 | 0.009 | 0.007 |
| ***missid*** | 0.004 | 0.006 | 0.005 | 0.003 | 0.005 | 0.004 | 0.001 | 0.002 | 0.001 | 0.002 | 0.008 | 0.005 | 0.002 | 0.007 | 0.004 | 0.003 | 0.004 | 0.003 |

*Demographic parameters inferred (in relation to Ne) using dadi with their confidence intervals for each Scenario.

**Table S2**

| Scenario | Simulated params | M1 vs M10 | M2 vs M20 | M1 vs M2 | M10 vs M20 | M20 vs M30 |
|---|---|---|---|---|---|---|
| 1 | m = 0, Sb = 0 and pc = 0 | 0.1239 | 0.4378 | 0.0000 | 0.0000 | 0.0132 |
| 2 | m = 0, Sb = 0 and pc = 0.05 | 0.3873 | 0.5393 | 0.0000 | 0.0000 | 0.0496 |
| 3 | m = 0, Sb = 0 and pc = 0.25 | 0.0838 | 0.4545 | 0.0095 | 0.0027 | 0.3323 |
| 4 | m = 0, Sb = 1 and pc = 0 | 0.5834 | 0.2364 | 0.0000 | 0.0000 | 0.0621 |
| 5 | m = 0, Sb = 1 and pc = 0.05 | 0.3288 | 0.2887 | 0.0000 | 0.0000 | 0.0450 |
| 6 | m = 0, Sb = 1 and pc = 0.25 | 0.1355 | 0.4304 | 0.0000 | 0.0000 | 0.4976 |
| 7 | m = 0, Sb = 10 and pc = 0 | 0.0187 | 0.2813 | 0.0000 | 0.0000 | 0.0001 |
| 8 | m = 0, Sb = 10 and pc = 0.05 | 0.1261 | 0.0718 | 0.0000 | 0.0000 | 0.0131 |
| 9 | m = 0, Sb = 10 and pc = 0.25 | 0.1302 | 0.3122 | 0.0017 | 0.0008 | 0.7474 |
| 10 | m = 0, Sb = 100 and pc = 0 | 0.0672 | 0.3442 | 0.0000 | 0.0000 | 0.0081 |
| 11 | m = 0, Sb = 100 and pc = 0.05 | 0.0105 | 0.4456 | 0.0000 | 0.0000 | 0.0000 |
| 12 | m = 0, Sb = 100 and pc = 0.25 | 0.6238 | 0.5614 | 0.0000 | 0.0000 | 0.1184 |
| 13 | m = 0.01, Sb = 0 and pc = 0 | 0.1921 | 0.3362 | 0.0000 | 0.0000 | 0.1599 |
| 14 | m = 0.01, Sb = 0 and pc = 0.05 | 0.4746 | 0.2638 | 0.0000 | 0.0000 | 0.2097 |
| 15 | m = 0.01, Sb = 0 and pc = 0.25 | 0.3522 | 0.4332 | 0.0001 | 0.0000 | 0.5744 |
| 16 | m = 0.01, Sb = 1 and pc = 0 | 0.3699 | 0.5464 | 0.0001 | 0.0001 | 0.3608 |
| 17 | m = 0.01, Sb = 1 and pc = 0.05 | 0.2405 | 0.2857 | 0.0000 | 0.0000 | 0.0970 |
| 18 | m = 0.01, Sb = 1 and pc = 0.25 | 0.0519 | 0.5359 | 0.0008 | 0.0001 | 0.2275 |
| 19 | m = 0.01, Sb = 10 and pc = 0 | 0.5686 | 0.5182 | 0.0000 | 0.0000 | 0.7076 |
| 20 | m = 0.01, Sb = 10 and pc = 0.05 | 0.4720 | 0.1078 | 0.0000 | 0.0000 | 0.3177 |
| 21 | m = 0.01, Sb = 10 and pc = 0.25 | 0.1691 | 0.3931 | 0.0073 | 0.0044 | 0.7500 |
| 22 | m = 0.01, Sb = 100 and pc = 0 | 0.4348 | 0.4815 | 0.0000 | 0.0000 | 0.3658 |
| 23 | m = 0.01, Sb = 100 and pc = 0.05 | 0.3306 | 0.4488 | 0.0000 | 0.0000 | 0.1923 |
| 24 | m = 0.01, Sb = 100 and pc = 0.25 | 0.4978 | 0.3463 | 0.0000 | 0.0000 | 0.1555 |

**Significance**

| Scenario | Simulated params | M1 vs M10 | M2 vs M20 | M1 vs M2 | M10 vs M20 | M20 vs M30 |
|---|---|---|---|---|---|---|
| 1 | m = 0, Sb = 0 and pc = 0 | FALSE | FALSE | TRUE | TRUE | TRUE |
| 2 | m = 0, Sb = 0 and pc = 0.05 | FALSE | FALSE | TRUE | TRUE | TRUE |
| 3 | m = 0, Sb = 0 and pc = 0.25 | FALSE | FALSE | TRUE | TRUE | FALSE |
| 4 | m = 0, Sb = 1 and pc = 0 | FALSE | FALSE | TRUE | TRUE | FALSE |
| 5 | m = 0, Sb = 1 and pc = 0.05 | FALSE | FALSE | TRUE | TRUE | TRUE |
| 6 | m = 0, Sb = 1 and pc = 0.25 | FALSE | FALSE | TRUE | TRUE | FALSE |
| 7 | m = 0, Sb = 10 and pc = 0 | TRUE | FALSE | TRUE | TRUE | TRUE |
| 8 | m = 0, Sb = 10 and pc = 0.05 | FALSE | FALSE | TRUE | TRUE | TRUE |
| 9 | m = 0, Sb = 10 and pc = 0.25 | FALSE | FALSE | TRUE | TRUE | FALSE |
| 10 | m = 0, Sb = 100 and pc = 0 | FALSE | FALSE | TRUE | TRUE | TRUE |
| 11 | m = 0, Sb = 100 and pc = 0.05 | TRUE | FALSE | TRUE | TRUE | TRUE |
| 12 | m = 0, Sb = 100 and pc = 0.25 | FALSE | FALSE | TRUE | TRUE | FALSE |
| 13 | m = 0.01, Sb = 0 and pc = 0 | FALSE | FALSE | TRUE | TRUE | FALSE |
| 14 | m = 0.01, Sb = 0 and pc = 0.05 | FALSE | FALSE | TRUE | TRUE | FALSE |
| 15 | m = 0.01, Sb = 0 and pc = 0.25 | FALSE | FALSE | TRUE | TRUE | FALSE |
| 16 | m = 0.01, Sb = 1 and pc = 0 | FALSE | FALSE | TRUE | TRUE | FALSE |
| 17 | m = 0.01, Sb = 1 and pc = 0.05 | FALSE | FALSE | TRUE | TRUE | FALSE |
| 18 | m = 0.01, Sb = 1 and pc = 0.25 | FALSE | FALSE | TRUE | TRUE | FALSE |
| 19 | m = 0.01, Sb = 10 and pc = 0 | FALSE | FALSE | TRUE | TRUE | FALSE |
| 20 | m = 0.01, Sb = 10 and pc = 0.05 | FALSE | FALSE | TRUE | TRUE | FALSE |
| 21 | m = 0.01, Sb = 10 and pc = 0.25 | FALSE | FALSE | TRUE | TRUE | FALSE |
| 22 | m = 0.01, Sb = 100 and pc = 0 | FALSE | FALSE | TRUE | TRUE | FALSE |
| 23 | m = 0.01, Sb = 100 and pc = 0.05 | FALSE | FALSE | TRUE | TRUE | FALSE |
| 24 | m = 0.01, Sb = 100 and pc = 0.25 | FALSE | FALSE | TRUE | TRUE | FALSE |

**Table S3**

| Inferred Sb | Domestic | Wild | True parameters | Scenario |
|---|---|---|---|---|
| **0** | **7** | **0** | S1M=0, Sb=0, pc=0 | 1 |
| 1 | 56 | 62 | | |
| 10 | 37 | 38 | | |
| 100 | 0 | 0 | | |
| **0** | **7** | **1** | M=0, Sb=0, pc=0.05 | 2 |
| 1 | 55 | 62 | | |
| 10 | 35 | 35 | | |
| 100 | 3 | 2 | | |
| **0** | **32** | **6** | M=0, Sb=0, pc=0.25 | 3 |
| 1 | 58 | 88 | | |
| 10 | 3 | 5 | | |
| 100 | 7 | 1 | | |
| 0 | 0 | 0 | M=0, Sb=1, pc=0 | 4 |
| **1** | **99** | **99** | | |
| 10 | 1 | 1 | | |
| 100 | 0 | 0 | | |
| 0 | 11 | 0 | M=0, Sb=1, pc=0.05 | 5 |
| **1** | **88** | **99** | | |
| 10 | 1 | 1 | | |
| 100 | 0 | 0 | | |
| 0 | 27 | 0 | M=0, Sb=1, pc=0.25 | 6 |
| **1** | **73** | **100** | | |
| 10 | 0 | 0 | | |
| 100 | 0 | 0 | | |
| 0 | 13 | 0 | M=0, Sb=10, pc=0 | 7 |
| 1 | 77 | 90 | | |
| **10** | **10** | **10** | | |
| 100 | 0 | 0 | | |
| 0 | 2 | 0 | M=0, Sb=10, pc=0.05 | 8 |
| 1 | 76 | 77 | | |
| **10** | **21** | **22** | | |
| 100 | 1 | 1 | | |
| 0 | 18 | 0 | M=0, Sb=10, pc=0.25 | 9 |
| 1 | 81 | 99 | | |
| **10** | **1** | **1** | | |
| 100 | 0 | 0 | | |
| 0 | 0 | 0 | M=0, Sb=100, pc=0 | 10 |
| 1 | 0 | 0 | | |
| 10 | 46 | 46 | | |
| **100** | **54** | **54** | | |
| 0 | 0 | 0 | M=0, Sb=100, pc=0.05 | 11 |
| 1 | 1 | 1 | | |
| 10 | 70 | 70 | | |
| **100** | **29** | **29** | | |
| 0 | 0 | 0 | M=0, Sb=100, pc=0.25 | 12 |
| 1 | 67 | 67 | | |
| 10 | 17 | 17 | | |
| **100** | **16** | **16** | | |

| Inferred Sb | Domestic | Wild | True parameters | Scenario |
|---|---|---|---|---|
| **0** | **4** | **0** | M=0.01, Sb=0, pc=0 | 13 |
| 1 | 48 | 50 | | |
| 10 | 48 | 50 | | |
| 100 | 0 | 0 | | |
| **0** | **12** | **0** | M=0.01, Sb=0, pc=0.05 | 14 |
| 1 | 35 | 40 | | |
| 10 | 45 | 52 | | |
| 100 | 8 | 8 | | |
| **0** | **8** | **5** | M=0.01, Sb=0, pc=0.25 | 15 |
| 1 | 84 | 92 | | |
| 10 | 1 | 1 | | |
| 100 | 7 | 2 | | |
| 0 | 0 | 0 | M=0.01, Sb=1, pc=0 | 16 |
| **1** | **99** | **99** | | |
| 10 | 1 | 1 | | |
| 100 | 0 | 0 | | |
| 0 | 5 | 8 | M=0.01, Sb=1, pc=0.05 | 17 |
| **1** | **62** | **66** | | |
| 10 | 18 | 19 | | |
| 100 | 15 | 7 | | |
| 0 | 24 | 0 | M=0.01, Sb=1, pc=0.25 | 18 |
| **1** | **74** | **0** | | |
| 10 | 0 | 98 | | |
| 100 | 2 | 0 | | |
| 0 | 1 | 0 | M=0.01, Sb=10, pc=0 | 19 |
| 1 | 73 | 73 | | |
| **10** | **26** | **27** | | |
| 100 | 0 | 0 | | |
| 0 | 2 | 3 | M=0.01, Sb=10, pc=0.05 | 20 |
| 1 | 0 | 0 | | |
| **10** | **90** | **92** | | |
| 100 | 8 | 5 | | |
| 0 | 0 | 3 | M=0.01, Sb=10, pc=0.25 | 21 |
| 1 | 97 | 97 | | |
| **10** | **0** | **0** | | |
| 100 | 3 | 0 | | |
| 0 | 0 | 0 | M=0.01, Sb=100, pc=0 | 22 |
| 1 | 12 | 12 | | |
| 10 | 73 | 73 | | |
| **100** | **15** | **15** | | |
| 0 | 0 | 0 | M=0.01, Sb=100, pc=0.05 | 23 |
| 1 | 0 | 0 | | |
| 10 | 0 | 0 | | |
| **100** | **100** | **100** | | |
| 0 | 0 | 0 | M=0.01, Sb=100, pc=0.25 | 24 |
| 1 | 36 | 36 | | |
| 10 | 20 | 20 | | |
| **100** | **44** | **44** | | |
